# Force sensing GPR133 is essential for normal balance and modulates vestibular hair cell membrane excitability via Gi signaling and CNGA3 coupling

**DOI:** 10.1101/2023.10.24.563882

**Authors:** Zhao Yang, Shu-Hua Zhou, Ming-Wei Wang, Yu-Qi Ping, Xiao-Long Fu, Ru-Jia Zhao, Peng Xiao, Yan Lu, Qi-Yue Zhang, Zhi-Chen Song, Yue-Tong Xi, Hui Lin, Yuan Zheng, Wei Qin, Fan Yi, Xiao Yu, Ren-Jie Chai, Jin-Peng Sun

## Abstract

The maintenance of normal balance sensing is a fundamental prerequisite for virtually every activity of daily life. As one set most important balance information collectors, vestibular hair cells convert mechanical stimuli from head movement into electrical signals through a mechanoelectrical transduction (MET) process. The molecular mechanism underlying equilibrioception and MET in vestibular hair cells is not well understood but is generally believed to be mediated by ion channels. However, whether these procedures are also mediated by other receptors, such as GPCRs, which are known to govern light, odorant and taste sensing, is not known. Here, by screening the expression of force-sensitive adhesion GPCRs in vestibular hair cells and phenotype profiling in animal models, we identified that a seven-transmembrane receptor, GPR133, was able to sense force in utricle hair cells and was required for maintenance of normal equilibrioception. Notably, GPR133 converted mechanical stimuli into changes in intracellular cAMP levels through Gi engagement and then modulated plasma membrane excitability and mediated MET by coupling to CNGA3 activity changes in approximately 30% of GPR133-expressing utricle hair cells. GPR133-mediated MET and its coupling with CNGA3 were recapitulated by an in vitro reconstitution system. Further chemical labeling, mass spectrometry and cryo-EM analysis provided potential structural information on force-induced GPR133 activation and Gi3 engagement. Collectively, our findings reveal the essential role of GPR133 in the maintenance of normal equilibrioception and suggest that GPCR family members can participate in the MET process in utricle hair cells through modulation of intracellular second messenger levels and ion channel coupling.

## Introduction

Balance sensing or equilibrioception, which ensures that we maintain orientation and stabilize visual input in three-dimensional space, is a fundamental prerequisite for virtually every activity of daily life. Our nervous system integrates balance information from three primary sensory systems: the vision, somatosensory and vestibular systems^1,2^. The vestibular system includes two otolith organs (utricle and saccule) and three semicircular canals with ampullae, which detect linear acceleration and rotational movement, respectively. Hair cells located at the vestibular sensory epithelium convert mechanical input from head movement into electrical signals through a mechanoelectrical transduction (MET) process, which is mediated by the directional deflection of the stereocilia (the hair bundles) sitting at the cuticular plate (the apical surface) of hair cells^3^. Although studies have shown that the regular distribution of these hair cells plays important roles in equilibrioception, the cellular mechanisms remain largely elusive. Clinical studies and animal-related research have revealed two categories of causal genes for vestibular dysfunction. The first gene group mainly modulates the functional homeostasis of the vestibular sensory epithelium; for instance, Ush1c affects the morphogenesis of stereocilia, Cyp26b1 regulates the zonal development of vestibular organs, and Pendrin maintains the electrolyte balance of the endolymph^4–6^. The second group of balance-regulating molecules is directly involved in the MET process, such as proteins expressed at the tip of the stereocilia, including PCDH15, TMHS, TMIE, and TMC1/2^7–9^. In particular, TMC1/2 has been suggested to be a pore-forming subunit of the MET channel, and double knockout of *Tmc1* and *Tmc2* in mice causes vestibular dysfunction^10,11^. However, direct evidence of force-induced TMC1/2 current has not been established in either an *in vitro* recombinant system or *in vivo*. Recently, another mechanosensory channel, Piezo2, was found to be expressed at the cuticular plate of hair cells, where it regulates functions distinct from the MET channels housed in stereocilia^12^.

In addition to ion channels, certain membrane receptors belonging to the family of G protein-coupled receptors (GPCRs), which are the most common drug targets, are able to sense mechanical forces^13–19^. For example, several adhesion GPCRs (aGPCRs), which possess large and multidomain N-termini that allow receptors to interact with extracellular matrices, could respond to mechanical force stimulation^18,20–22^ (Figure S1A). Ion channels and GPCRs are two classes of sensory receptors. Notably, temperature and touch are sensed by TRP channels and piezo channels, respectively^23,24^. In contrast, vison and olfaction are mainly mediated by GPCRs, which convert light or odor stimuli into electrical signals through coupling to cyclic nucleotide-gated (CNG) channels^25–28^. Both GPCR family members and ion channels participate in distinct taste sensation^29,30^. The downstream signaling and cellular outputs differ between GPCRs and ion channels. Whereas sensory ion channels directly mediate ion permeability, GPCRs regulate sensory signals by controlling the intracellular concentrations of secondary messengers (cAMP, cGMP, Ca^2+^, etc.). Although investigations into the functions of channels in equilibrioception have emerged recently, there is no report that GPCRs are able to sense mechanical force and transduce it into electrical signals or participate in balance functions.

Here, we screened the mechanosensitive potential of adhesion GPCRs (GPCRs with force-sensing potential) in utricle hair cells via expression level analyses, cellular assays and animal models. We identified one particular receptor, GPR133, as a mechanosensor that is essential for equilibrioception. Importantly, systemic deletion of *Gpr133* or conditional deficiency induced by *Pou4f3-CreER*^+/-^*Gpr133*^fl/fl^ caused balance dysfunctions and impaired equilibrioception. Moreover, we obtained direct evidence that GPR133 converts mechanical stimulation into electrical signals by downregulating cAMP levels in utricle hair cells using recombinant systems. Force sensing by GPR133 is coupled to CNGA3 and modulates plasma membrane excitability. We therefore report that a force-sensing GPCR is able to participate in MET and is essential for equilibrioception.

## Results

### Deficiency of the mechanosensor GPR133 impaired balance but not hearing function

The human aGPCR family consists of 33 members (30 in mice), and several members have been reported to be activated by mechanical forces (Figure S1A)^17,18,21,31–33^. Single-cell RNA sequencing (scRNA-seq) data indicated that twelve aGPCRs were expressed in more than 20% of utricle hair cells of mice (Figure 1A). We then investigated the expression patterns of these twelve receptors in *Slc26a4*-deficient mice, which show vestibulogenic balance disorder and are widely used models for enlargement of the vestibular aqueduct (EVA)^34^. Notably, the expression levels of five aGPCRs, including GPR133, GPR123, CELSR1, GPR97 and GPR126, were significantly changed in the vestibular organs of the *Slc26a4*-deficient mouse model, which may correlate with balance function in the vestibular system (Figure 1B).

**Figure 1.**
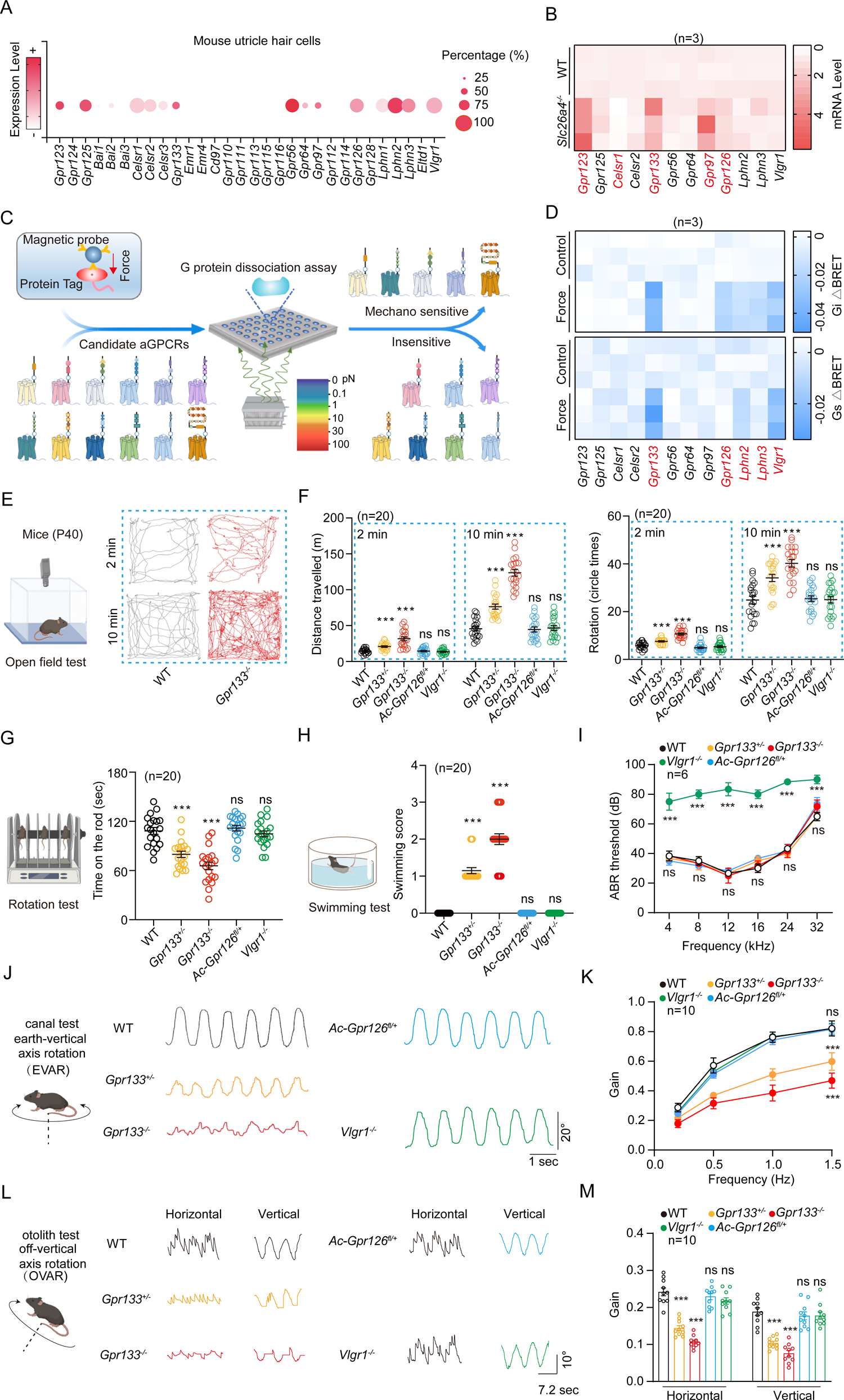
Deficiency of mechano-sensing GPR133 impairs balance. **(A)** Expression profiles of 30 adhesion GPCR genes in mouse utricle hair cells. Single cell RNA-seq datasets of utricle hair cells are from GSE71982. The size of each circle represents the percentage of hair cells where the gene is detected, and the color intensity represents the average mRNA expression level of the gene. **(B)** mRNA levels of 12 aGPCR genes in the utricle isolated from WT or *Slc26a4^-/-^* mice (n=3). **(C)** Schematic representation of the strategy used to screen the mechano-sensing aGPCR. **(D)** Summary of the force-induced Gi (top panel) and Gs (bottom panel) activation of 12 aGPCRs. A force of 3 pN was applied to the receptors and the Gi or Gs activation was measured by BRET assay, which was presented as a heatmap (n=3). **(E)** Schematic view (left) and representative tracks (right) of WT and *Gpr133^-/-^* mice in open-field test during 2 min or 10 min tracking period. **(F)** Quantification of the travelling (left) and circling activity (right) of WT, *Gpr133^-/-^*, *Gpr133^+/-^*, *Vlgr1^-/-^*, and *Atoh1-cre*^+/-^*Gpr126*^fl/+^ (referred to as *Ac-Gpr126^+/-^*) mice in open-field test (n=20 mice per group). **(G)** Schematic view (left) and quantification (right) of time on the rotating rod of WT, *Gpr133^-/-^*, *Gpr133^+/-^*, *Vlgr1^-/-^*, and *Ac-Gpr126^+/-^* mice in rotation test (n=20 mice per group). **(H)** Schematic view (left) and quantification (right) of swimming scores of WT, *Gpr133^-/-^*, *Gpr133^+/-^*, *Vlgr1^-/-^*, and *Ac-Gpr126^+/-^* mice in forced swimming test (n=20 mice per group). **(I)** ABR responses obtained for WT, *Gpr133^-/-^*, *Gpr133^+/-^*, *Vlgr1^-/-^*, and *Ac-Gpr126^+/-^* mice (n=6 mice per group). **(J)** Schematic view (left) and representative recording curves (right) of the VOR response to earth-vertical axis rotation (0.25-1.5 Hz, 40°/s peak velocity sinusoidal, whole-body passive rotation). **(K)** Quantification of the VOR gain response of WT, *Gpr133^-/-^*, *Gpr133^+/-^*, *Vlgr1^-/-^*, and *Ac-Gpr126^+/-^* mice to earth-vertical axis rotation (n=10 mice per group). **(L)** Schematic view (left) and representative recording curves (right) of the VOR response to off-vertical axis rotation (50°/s, whole-body passive rotation). **(M)** Quantification of the VOR gain response of WT, *Gpr133^-/-^*, *Gpr133^+/-^*, *Vlgr1^-/-^*, and *Ac-Gpr126^+/-^* mice to off-vertical axis rotation (n=5 mice per group). (**F-I, K, M**) **P < 0.01; ***P < 0.001; ns, no significant difference. Gene knockout mice compared with WT mice. The bars indicate mean ± SEM values. All data were statistically analyzed using one-way ANOVA with Dunnett’s post hoc test.

We then established a high-throughput mechanical stimulation assay to examine the mechanosensing ability of these twelve aGPCRs that were enriched in the vestibular system using a magnetic tweezer system connected with a GPCR biosensor platform. Briefly, tension forces were applied to paramagnetic beads coated with anti-Flag M2 antibody or with polylysine using a magnetic system, and force-induced G protein activation was examined using a bioluminescence resonance energy transfer (BRET) assay in HEK293 cells transfected with plasmids encoding N-terminal Flag-tagged GPCR and G protein biosensors (Figure 1C)^35,36^. In response to magnetic stimulation, the tension force was produced by magnetic beads attached to the N-terminus of selected aGPCRs, and the activation of Gs or Gi signaling downstream of mechanosensitive GPCRs was measured by BRET signals. The Gs/Gi signal was selected as a readout because recent studies have indicated critical roles of Gi protein and cAMP levels in the regulation of stereocilium development, vestibular function and hair cell mechanotransduction^37–39^. Using this system, we revealed that 5 receptors (GPR133, GPR126, LPHN2, LPHN3 and VLGR1) activated Gi signaling in response to force stimulation, whereas 3 receptors (GPR133, LPHN2 and VLGR1) activated Gs signaling in response to force application (Figure 1D). The specific changes in expression levels in the vestibular system of mouse models exhibiting balance disorders and the mechanosensitive capability of GPR133 and GPR126 collectively prompted us to explore the potential functions of these two receptors in balance. In addition, we included VLGR1 for balance phenotype analysis because VLGR1 was able to sense force, and its mutants were highly associated with deafness, a phenotype tightly connected with balance^40,41^. We then investigated whether these mechanosensitive aGPCRs are essential for equilibrioception by comparing *Gpr133*^+/-^ mice, *Gpr133*^-/-^ mice, *Atoh1-cre*^+/-^*Gpr126*^fl/+^ mice and *Vlgr1*^-/-^ (*Vlgr1*/del7TM) mice with their wild-type (WT) littermates in different equilibration-related tests (Figures S1B-S1J). To avoid potential neonatal death, we generated *Atoh1-cre*^+/-^*Gpr126*^fl/+^ mice (Figures S1H-S1J). *Atoh1* is highly enriched in nascent cochlear and vestibular hair cells and is commonly used as a marker of hair cells^42,43^. The scRNA-seq data suggested that approximately 85% of *Gpr126*-expressing utricle hair cells were *Atoh1*-positive. The deficiency of GPR133, GPR126, or VLGR1 in the mutant mice was verified by genotyping PCR (Figures S1C, S1F, and S1I) and western blotting analysis (Figures S1D, S1G, and S1J). All the above mice were viable, fertile and had normal body weights under normal chow diet feeding (Figure S 2A). The vestibular and auditory behaviors of the mice at P40 were assessed. Notably, the *Gpr133*^+/-^ mice exhibited increased circling behavior compared with that of their WT littermates in both the 2 min time frame (7.6 ± 0.3 circles/7.8 ± 0.8 circles vs. 5.8 ± 0.3 circles) and the 10 min time frame (34.1 ± 1.5 circles/38.6 ± 2.3 circles vs. 24.9 ± 1.6 circles) (Figures 1E-1F). The *Gpr133*^-/-^ genotype aggravated these phenotypes in both the 2 min (10.7 ± 0.4 circles vs. 5.8 ± 0.3 circles) and 10 min time frames (40.2 ± 1.6 circles vs. 24.9 ± 1.6 circles). In contrast, the *Atoh1-cre*^+/-^*Gpr126*^fl/+^ mice and *Vlgr1*^-/-^ mice showed circling behaviors similar to those of their WT littermates (Figure 1F). The vestibular function of the mice was further examined using a rotarod apparatus, and the holding time of the tested mice on the rotating rod before falling off was recorded. The *Gpr133*^+/-^ and *Gpr133*^-/-^ mice showed approximately 25% and 40% shorter durations in the rotarod test than the WT mice, respectively (Figure 1G). These results implied that *Gpr133-deficient* mice had severe balance defects. Consistently, the *Gpr133*^+/-^ and *Gpr133*^-/-^ mice showed impaired performance in the forced swim test, with swimming scores of 1.2 ± 0.1 and 2.0 ± 0.2, respectively (the 0–3 scoring system was employed, and WT mice had scores of 0^44^) (Figure 1H). In contrast, the *Atoh1-cre*^+/-^*Gpr126*^fl/+^ mice and *Vlgr1*^-/-^ mice exhibited no significant difference compared with the WT mice in rotarod apparatus rotating or forced swimming tests (Figures 1G-1H). Importantly, the balance defect of *Gpr133*^-/-^ mice was not due to cerebellum defects, since the cerebella of the *Gpr133^-/-^* mice had normal morphologies and Purkinje cell numbers, showing no significant differences compared with those of WT mice (Figures S2B-S2D). Moreover, the threshold of auditory brainstem responses (ABR) of the *Gpr133-*deficient mice was comparable with that of WT mice (Figure 1I, Figure S2E). The outer hair cells of *Gpr133-*deficient mice showed normal function, as indicated by distortion product otoacoustic emission (DPOAE) measurements (Figure S2F). As a control, *Vlgr1*^-/-^ mice showed abnormalities in both the ABR and DPOAE tests (Figure 1I, Figures S2E-S2F). Finally, *Gpr133*^+/-^ mice and *Gpr133*^-/-^ mice, but not *Atoh1-cre*^+/-^*Gpr126*^fl/+^ or *Vlgr1*^-/-^ mice, exhibited significantly impaired vestibulo-ocular reflexes (VORs) in response to both earth-vertical and off-vertical axis rotations compared with those of WT mice (Figures 1J-1M).

Taken together, these results indicated that mechanosensitive GPR133 played an important role in regulating equilibrioception, and deletion of *Gpr133* in mice significantly impaired balance but not hearing function.

### Expression of GPR133 in utricle hair cells

The mammalian vestibular system includes three semicircular canals and two macular organs. The two macular organs, utricle and saccule, have been proposed to detect linear acceleration and head tilt with respect to gravity. Both the utricle and saccule have two anatomical zones, namely, the central striola (S) region and the peripheral extrastriola (ES) region, which show distinct features such as otoconia size and hair cell morphology (Figure 2A)^45,46^. To specifically label GPR133 expression *in vivo* and circumvent the potential nonspecificity of antibody staining, we generated transgenic KI mice by inserting a GFP tag into the C-terminus of GPR133 (Figures S3A-3B). Whole-mount utricles derived from KI mice were immunostained with calbindin, a selective S region hair cell marker. The results suggested that GPR133 was more enriched in the S region (approximately 70%) than in the lateral ES (LES) region (approximately 30%) of the utricle in WT mice (Figures 2B-2D). The expression pattern of GPR133 was further supported by RNAscope *in situ* hybridization and immunostaining with GPR133 antibody (Figure 2C, Figure S3C). The specificity of the GPR133 antibody was examined by both immunostaining of utricles derived from *Gpr133*^-/-^ mice and western blotting of GPR133-overexpressing HEK293 cells (Figures S3C-S3D). GPR133 expression was restricted to hair cells on both sides of the line of polarity reversal (LPR) but not supporting cells, as evidenced by its coimmunostaining with oncomodulin (OCM, a hair cell marker in the S region) and osteopontin (OPN, a hair cell marker in the ES region) but not with SOX2 (a supporting cell marker) (Figure 2E). Furthermore, section staining of the utricle sensory epithelium revealed that GPR133 was mostly localized at the cuticular plate and cell surface of hair cells (Figure 2E). In particular, GPR133 localization was also found at the basal membrane connecting with the supporting cells or afferent neurons. Intriguingly, GPR133 expression was not detected within the stereocilia (Figure S3E).

**Figure 2.**
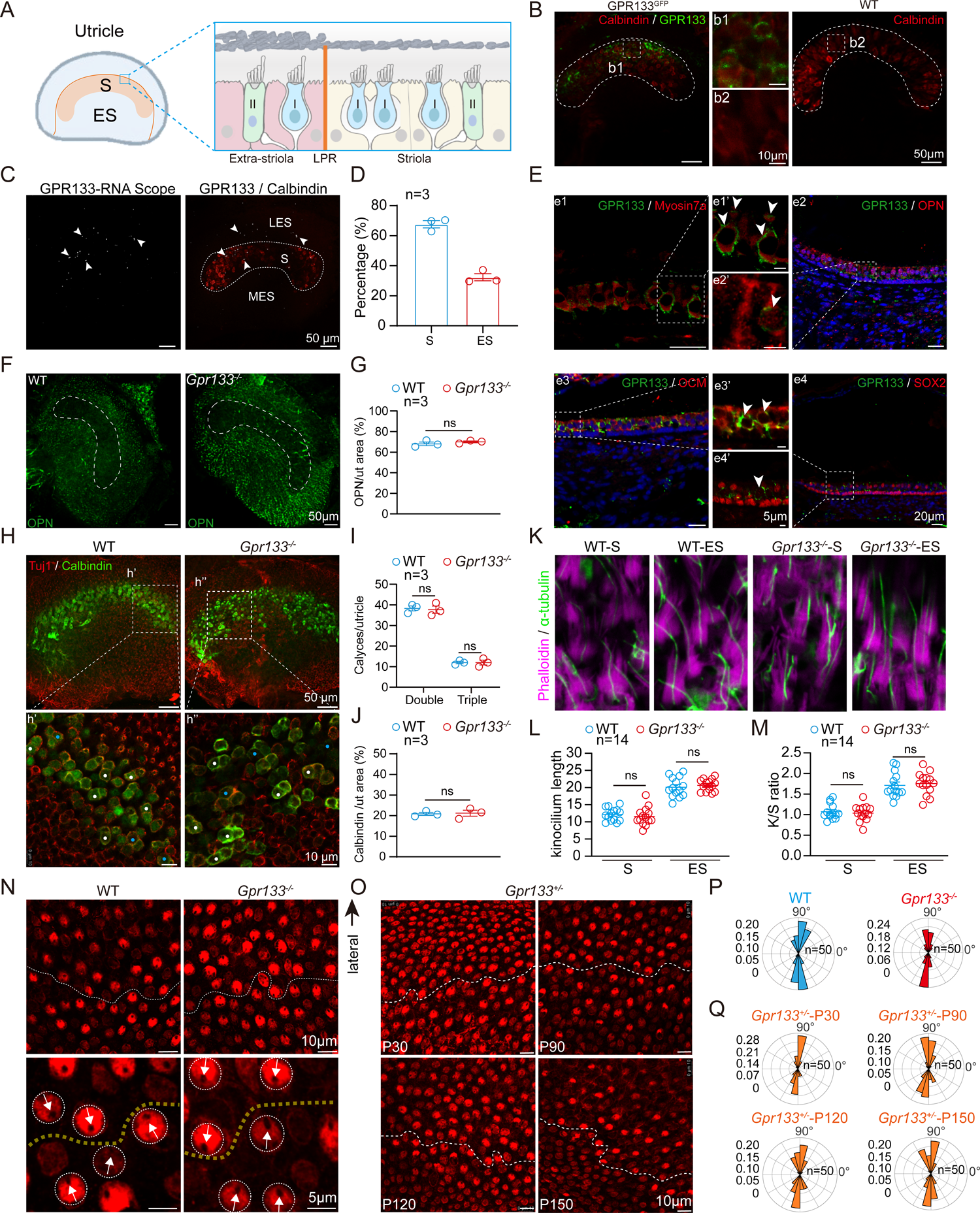
GPR133 is dispensable for zonal development and hair cell polarity in utricle. **(A)** Schematic representation of the section view of the utricle across the striola (S) and extra-striola (ES) region. The LPR line is depicted in orange. **(B)** Immunostaining of calbindin (red) in utricle whole mounts derived from *GPR133^GFP^* (b1) and WT (b2) mice at P40. Enlarged images show the expression of GPR133 (green) at the surface of hair cells in S regions (dotted outline) (n = 3 mice per group). Scale bars: 50 μm and 10 μm for low and high magnification view, respectively. **(C)** Representative images of whole-mount RNAscope *in situ* hybridization of GPR133 (white) combined with immunostaining of calbindin (red) in the utricle of P40 mice (n = 3 mice). Arrows indicate GPR133 staining in both S (dotted outline) and ES regions. Scale bar, 50 μm. **(D)** Quantitative analysis of the GPR133 expression in S and ES regions. Data are correlated to Figure 2B-2C (n = 3 mice). **(E)** Co-immunostaining of GPR133 (green) with myosin7a (e1-e1’, red, hair cell marker), osteopontin (e2-e2’, red, hair cell marker in ES region) oncomodulin (e3-e3’, red, hair cell marker in S region), or Sox2 (e4-e4’, red, supporting cell marker) in utricle sections derived from WT mice (n = 3 mice per group). GPR133 expression is restricted to hair cells at both sides of LPR but not supporting cells. Scale bars: 20 μm for e1, e2, e3, e4; 5 μm for e1’, e2’, e3’, e4’. **(F-G)** Immunostaining (**F**) and quantitative analysis (**G**) of osteopontin expression in utricle whole mounts derived from WT and *GPR133^-/-^* mice (n = 3 mice per group). Scale bar, 50 μm. **(H)** Co-immunostaining of Tuj1 (red) and calbindin (green) in utricle whole mounts derived from WT and *GPR133^-/-^* mice. Enlarged images show the double (white) and triple (blue) calyces encasing hair cells in S region. Scale bar: 50 μm and 10 μm for low (top panel) and high magnification (bottom panel) view, respectively **(I-J)** Quantification of the calyces numbers (**I**) and calbindin expression (**J**) in utricle whole mounts derived from WT and *GPR133^-/-^*mice. Data are correlated to Figure 2H (n = 3 mice per group). **(K)** Immunostaining of kinocilium (labeled with α-tubulin, green) and stereocilia (labeled with phalloidin, red) in utricle whole mounts derived from WT and *GPR133^-/-^*mice. Scale bar, 10 μm. **(L-M)** Quantification of the length of kinocilium (**L**) and the ratio of lengths of the kinocilium to tallest stereocilia (**M**) in utricle whole mounts derived from WT and *GPR133^-/-^* mice. Data are correlated to Figure 2K (n = 14 hair cells from 3 mice per group). **(N)** Immunostaining of βII-spectrin reveals the orientation of hair cells at LPR region of utricles derived from WT and *GPR133^-/-^* mice. The hair cell orientation is indicated by the position of the off-center fonticulus without signal. Scale bars: 10 μm and 5 μm for low (top panel) and high magnification (bottom panel) view, respectively. **(O)** The orientation of hair cells at LPR region of utricles derived from *GPR133^+/-^*mice at P30, P90, P120 and P150. Scale bar, 10 μm. **(P)** Circular histogram showing the frequency distribution of hair cell orientation at LPR region of utricles derived from WT and *GPR133^-/-^* mice (n=50 hair cells from 3 mice per group). The lateral side of utricle is defined as 90° (top direction in the histogram). Data are correlated to Figure 2N. **(Q)** Circular histogram showing the frequency distribution of hair cell orientation at LPR region of utricles derived from *GPR133^+/-^* mice at P30, P90, P120 and P150 (n=50 hair cells from 3 mice per group). Data are correlated to Figure 2O. (**G, I-J, L-M**) ns, no significant difference. *GPR133^-/-^* mice compared with WT mice. The bars indicate mean ± SEM values. All data were statistically analyzed using student’s t test (**G, J**) or two-way ANOVA with Dunnett’s post hoc test (**I, L-M**).

We next investigated the morphological and anatomical features of the utricle in *Gpr133*^-/-^ mice at P40, which showed significant balance deficits. The size of maculae and total hair cell numbers in the utricle of these *Gpr133*^-/-^ mice were comparable with those of WT mice (Figures S2F-S2G). Notably, the expression of OPN, the selective marker for ES hair cells, in *Gpr133*^-/-^ mice was comparable with that in their WT littermates (Figures 2F-2G). In addition, calbindin immunoreactivity and the numbers of complex calyces encasing two or three hair cells (identified by *Tuj1* staining), which are hallmarks of the S region, showed no significant differences in *Gpr133*^-/-^ mice compared to WT mice (Figures 2H-2J). These data suggested that the morphological integrity of the utricle and the regional identity of hair cells in *Gpr133*^-/-^ mice at P40 were normal compared with those in WT mice. This hypothesis was further supported by the evidence that the kinocilium length and the K/S ratio (the ratio of the length of the kinocilium (stained by α-tubulin) to that of the tallest stereocilium (stained by phalloidin)) in the S and ES regions of the *Gpr133^-/-^* utricle were comparable to those of their WT counterparts (Figures 2K-2M). Moreover, scanning electron microscopy (SEM) analysis revealed that *Gpr133* deficiency did not affect the size of the S region in the utricle or the density of otoconia in either the S or ES zone at P40 (Figures S2H-S2J).

The unique asymmetrical orientation, or polarity, of the stereocilia of hair cells determines the directional sensitivity and is pivotal for the normal functions of the vestibular system and the organ of Corti. Specifically, LPR divides the utricle into two regions (LES and S) with opposing stereocilia polarities, and loss of hair cell polarity in the LPR region is associated with vestibular deficits^47^. We therefore investigated the hair cell orientation in domains across the LPR in the utricular macula of WT and *Gpr133^-/-^* mice at P40 (Figure 2N). We found that the hair cells in the LPR region of the *Gpr133^-/-^* utricle displayed normal orientations, showing no significant difference compared with the WT utricle (Figures 2N and 2P). Moreover, although the *Gpr133^+/-^*mice exhibited balance defects, similar normal tissue organization and morphological integrity were observed for the *Gpr133^+/-^* mice at different postnatal time points, including P30, P90, P120 and P150 (Figures 2O and 2Q). These results collectively suggested that *Gpr133* might participate in an instantaneous physiological process in utricle hair cells in addition to potential developmental effects.

### Force-induced G protein activation and second messenger changes through activating GPR133

aGPCR members, such as GPR56, GPR64 and GPR126, are known to couple to multiple G protein subtypes for effective signaling^48–53^. In addition to Gi, previous studies have shown that *Stachel* sequence-derived peptides activate Gs signaling of GPR133, and we have recently solved the cryo-EM structure of the GPR133β-Gs complex^18,20,54^. Because the fluid force induced by head motion is one of the main physiological stimuli in the utricle, we next investigated G protein subtype activation patterns of GPR133 in response to force stimulation. We coated paramagnetic beads with either anti-Flag M2 antibody or anti-GPR133 (recognizing residues 479-501 at the N-terminus) antibody (hereafter referred to as Flag-M-beads or GPR133-M-beads, respectively) and then incubated them with GPR133-overexpressing HEK293 cells using polylysine-coated beads and pCDNA3.1 plasmid-transfected cells as a negative control (Figure 3A). We used the magnetic field to control the force administered to the GPR133 receptor, which was calculated according to the following formula: F_z_=(−1.2×10^-10^)*exp*(–104Z) (Figure 3B). We then employed force stimulation with a magnetic field and found that both the Gi and Gs pathways downstream of GPR133 were activated (Figures 3C-3D, Figures S4A-S4C). The EC50 values for force-induced GPR133 activation of Gs, Gi1, Gi2 and Gi3 signaling using Flag-M-beads were 2.79 ± 0.03, 2.73 ± 0.03, 2.49 ± 0.03 and 2.28 ± 0.04 pN, respectively (Figure 3D). In contrast, GPR133 could not translate the mechanical force into detectable activation of Go, Gq, G12 or β-arrestin signaling (Figures S4D-S4H).

**Figure 3.**
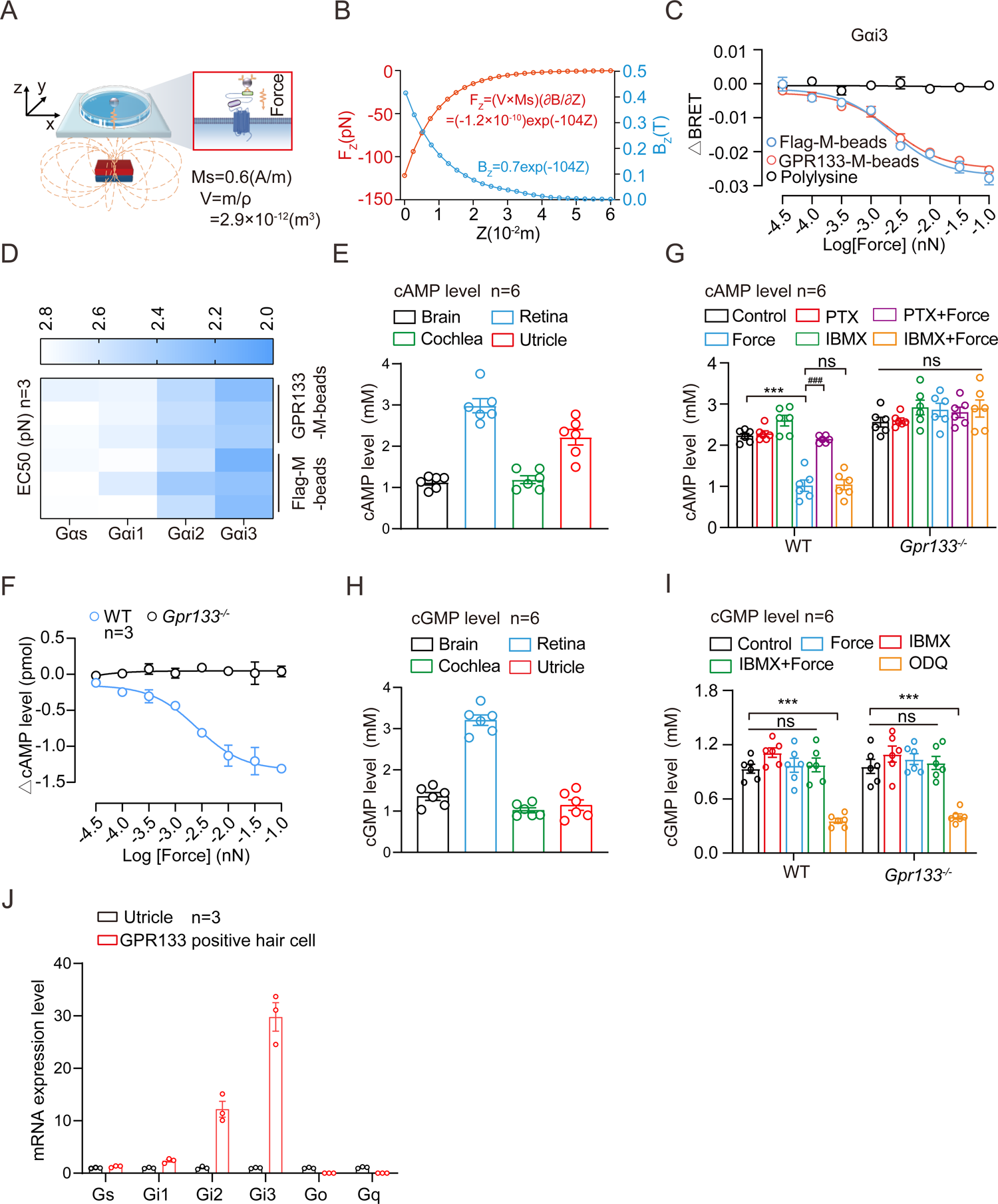
Force induces cAMP reduction in utricle hair cells through activating GPR133-Gi signaling. **(A)** Schematic representation of the force-induced GPR133 activation in HEK293 cells using a magnetic system. **(B)** Magnetic force (Fz) as a function of distance (Z) between the magnetic beads and the permanent magnet. **(C)** Dose-dependent Gi3 activation in GPR133-overexpressed HEK293 cells in response to force applied on paramagnetic beads coated with anti-Flag M2 or anti-GPR133 antibody. Polylysine-coated beads were used as a negative control (n=3). **(D)** Summary of the potency of force-induced Gs or Gi activation in GPR133-overexpressed HEK293 cells (n=3). Data are correlated to Figure S4. **(E)** Endogenous cAMP levels in different tissues including brain, retina, cochlea and utricle measured by ELISA kit (n=6). **(F)** Force-induced dose-dependent reduction of cAMP levels in the utricle explant derived from WT and *GPR133^-/-^* mice (n=3). **(G)** Effects of Gi inhibitor PTX or PDE inhibitor IBMX on the force-induced cAMP reduction in WT utricle explant (n=6). **(H)** Endogenous cGMP levels in different tissues including brain, retina, cochlea and utricle (n=6). **(I)** Effects of force on cGMP levels in WT utricle explant (n=6). The guanylyl cyclase inhibitor ODQ was used as a positive control. **(J)** mRNA levels of different G protein subtypes in the utricle and GPR133-positive utricle hair cells (n=3). (G) ***P < 0.001; ns, no significant difference. Force-stimulated utricle explant compared with control utricle explant. ^###^P < 0.001; ns, no significant difference. PTX- or IBMX-treated utricle explant compared with control utricle explant. (I) ***P < 0.001; ns, no significant difference. Force and/or reagents treated utricle explant compared with control utricle explant. The bars indicate mean ± SEM values. All data were statistically analyzed using one-way ANOVA with Dunnett’s post hoc test.

The Gs and Gi coupling property of mechanosensitive GPR133 suggested that it might regulate intracellular cAMP or cGMP levels in the utricle. We therefore incubated the utricles derived from both *Gpr133*^-/-^ mice and their WT littermates with GPR133-M-beads, which showed comparable potency and efficacy in activating Gs and Gi signaling downstream of GPR133 with the Flag-M-beads in a heterologous system (Figure 3C-3D, Figures S4A-S4C). Notably, the basal intracellular cAMP levels of the utricle and retina were approximately 2.22±0.19 mM and 2.98±0.17 mM, respectively, which were significantly higher than those of the brain or cochlea (approximately 1 mM) (Figure 3E). Administration of magnetic force to the utricle hair cells by GPR133-M-beads induced a dose-dependent decrease in the basal cAMP level in the utricle explant of WT mice with an EC50 of 2.57±0.11 pN but not in the *Gpr133*^-/-^ utricle explant (Figures 3F-3G). In contrast, the cGMP levels in both WT and *Gpr133*^-/-^ utricles showed no significant changes in response to force application (Figures 3H-3I). Notably, the force-induced cAMP decrease could be inhibited by PTX treatment but not by IBMX, suggesting that force modulated intracellular cAMP levels of GPR133-expressing hair cells in the utricle through GPR133-Gi signaling (Figure 3G). The force-induced cAMP decrease via GPR133 activation was consistent with the higher Gi expression levels than Gs levels in the utricle and GPR133-positive hair cells (Figure 3J).

### Activation of GPR133 by force modulates the excitability of utricle hair cells

The balance defects without abnormal utricle morphology in *Gpr133*^-/-^ mice at P40 suggested that GPR133 might participate in an instantaneous physiological process in utricle hair cells. We next investigated whether activation of GPR133 by force affects the excitability of utricle hair cells. The promoter region of GPR133 was cloned into AAV-ie vector to replace the original CAG promoter and direct the expression of GFP (AAV-ie-*Gpr133pr*-GFP), which enabled specific labeling of GPR133-expressing hair cells in the utricle, as verified by both single-cell RT‒PCR and fluorescent colocalization (Figure 4A, Figures S5A-S5B)^55^. Whole-cell patch-clamp recordings from GPR133-promoter labeled WT utricle hair cells were carried out in response to stimulation with 3 pN force using GPR133-M-beads. Importantly, 39% of GPR133-expressing hair cells were force sensitive (cluster I), as evidenced by an approximately 50% reduction in current amplitude in response to force stimulation, whereas the other 61% of GPR133-expressing hair cells were force insensitive (cluster II) (Figures 4B-4D, Figures S5C-S5D). None of the GPR133 promoter-labeled utricle hair cells derived from *Gpr133*^-/-^ mice (negative controls) were sensitive to force using GPR133-M beads (Figures 4B-4D).

**Figure 4.**
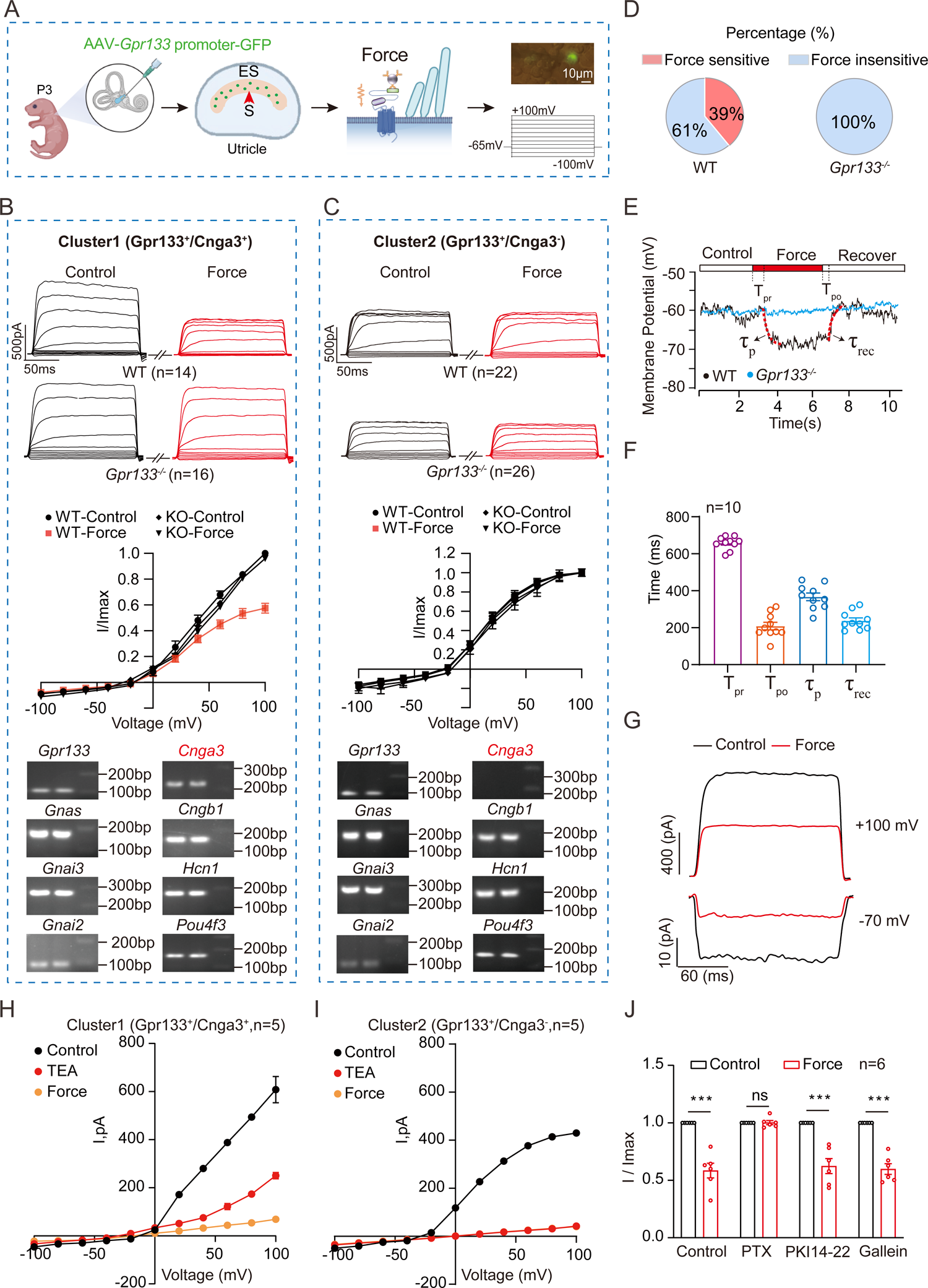
Force sensing by GPR133 modulates membrane excitability of utricle hair cells. **(A)** Schematic illustration of force application and patch-clamping on *Gpr133* promoter-labeled utricle hair cells. **(B-C)** Representative whole-cell currents (top panel) elicited by voltage steps between −100 mV and +100 mV and the corresponding I-V curves (middle panel) in *Gpr133* promoter-labeled cluster I (B) or cluster II (C) utricle hair cells derived from WT or *Gpr133^-/-^*mice in response to force (3 pN) stimulation. The expression of indicated genes in hair cells of both clusters were examined by single cell RT-PCR (bottom panel). n=14 for cluster I cells and n=22 for cluster II cells in WT mice. n=16 for cluster I cells and n=26 for cluster II cells in *Gpr133^-/-^* mice. Data are correlated to Figure S5C-S5D. **(D)** Quantification of force-sensitive and force-insensitive utricle hair cells derived from WT or *Gpr133^-/-^*mice. Data are correlated to Figure 4B-4C. **(E)** Representative force-induced membrane potential alteration of cluster I utricle hair cells derived from WT or *Gpr133^-/-^* mice when holding the cells at absolute zero current. The delayed time between force stimulation or force removal and the change of membrane potential was defined as T_pr_ and T_po_, respectively. The calculated time constant for response and the recovery time constant was denoted as τ_p_ and τ_rec_, respectively. **(F)** The calculated time parameters in force-stimulated membrane potential alteration (n=10). **(G)** Representative whole cell currents in cluster I hair cells derived from WT mice in response to force stimulation (3 pN) at a holding potential of +100 mV or −70 mV. **(H-I)** Effect of potassium channel blocker TEA on the whole-cell current in cluster I (**H**) and cluster II (**I**) hair cells derived from WT mice without or with force (3 pN) stimulation. Data are correlated to Figure S5J. **(J)** Effect of PTX, the PKA inhibitor PKI 14-22 or the Gβγ inhibitor galleon pre-treatment on the force-induced whole-cell current reduction in cluster I hair cells at a holding potential of +100 mV (n=6). (**J**) ***P<0.001; ns, no significant difference. Force-stimulated cluster I hair cells compared with control cluster I hair cells. The bars indicate mean ± SEM values. All data were statistically analyzed using two-way ANOVA with Dunnett’s post hoc test.

To assess the excitability of utricle hair cells in response to force stimulation administered on GPR133, we recorded the resting membrane potential of cluster I cells under a current clamp. When holding at 0 pA, repeatable force application induced dose-dependent hyperpolarization of the plasma membrane of cluster I utricle hair cells with an EC50 of 2.55 ± 0.51 pN, similar to those recorded in both the G protein activation assay in the heterologous system and the cAMP reduction in utricle explants (Figure 4E and Figures S5F-S5G). Notably, force application using Flag-M-beads on the utricle hair cells derived from the Flag-GPR133^GFP^ knock in mice also induced a dose-dependent hyperpolarization with an EC50 (2.47 ± 0.18 pN) similar to that recorded in WT utricle hair cells in response to GPR133-antibody conjugated M-beads, further supporting the specificity of force sensation by GPR133 (Figure S5H). The hair cell membrane potential was rapidly changed in response to force administration, with a delayed time of 658.0 ± 11.14 ms, and could be recovered after 3 s of force stimulation (recovery delayed by 208.9 ± 20.59 ms) (Figures 4E-4F). The response kinetics of the force-induced voltage change were best fitted with a mono-exponential equation. The calculated time constant for response (τ_p_) was 367.0 ±52.1 ms, while the recovery time constant (τ_rec_) was 237.4 ± 30.5 ms when the cells were held at absolute zero current (Figure 4F). Consistently, the current in cluster I cells was significantly decreased in response to force stimulation at a holding potential of either +100 mV or −70 mV (Figure 4G). Both force-induced membrane potential alteration and current reduction in response to GPR133-M-beads were abolished in GPR133-promoter labeled utricle hair cells from *Gpr133*^-/-^ mice (Figure 4E, Figure S5I).

Pronounced outward rectification currents and a reversed potential of approximately −10 mV were observed in both cluster I and cluster II cells (Figures 4B-4C). The current-voltage relationship was almost linear from −10 to 100 mV in cluster I cells (Figure 4B). In contrast, the current-voltage relationship was only almost linear from −10 to 50 mV in cluster II cells (Figure 4C). Interestingly, although the GPR133 promoter-labeled utricle hair cells derived from *Gpr133*^-/-^ mice showed no response to the force applied to GPR133-M-beads, these cells could still be divided into two clusters according to the differences in the current-voltage relationship, similar to WT mice (Figures 4B-4C). These results indicated that GPR133 deficiency does not directly affect the basal excitability of these hair cells. Importantly, including tetraethylammonium (TEA) in the extracellular solution decreased the whole-cell current in both cluster I and cluster II cells but showed no significant effect on the force-induced current alteration in cluster I cells (Figures 4H-4I, Figure S5J). These results indicated that potassium contributed to the excitability of GPR133-expressing hair cells, but the potassium-related excitability was not modulated by the force sensed by GPR133. Moreover, the force-induced current was completely blocked by pretreatment with PTX but was not affected by PKI 14-22 (PKA inhibitor) or galleon (Gβγ inhibitor) (Figure 4J). Collectively, these results suggest that the GPR133-regulated conversion of force stimulation into electronical signals in utricle hair cells is Gi-dependent and most likely potassium channel-independent.

### Functional coupling of GPR133 with CNGA3

We next looked for downstream effectors of GPR133-Gi-CAMP signaling in GPR133-expressing utricle hair cells for modulation of membrane excitability in response to mechanosensing of GPR133. The expression of cyclic nucleotide–activated ion channels, including cyclic nucleotide–gated (CNG) channels and hyperpolarization-activated cyclic nucleotide–modulated (HCN) channels, in utricle hair cells was examined. Only 5 members (CNGA1, CNGA3, CNGB1, HCN1 and HCN4) of these two channel families were detected in GPR133-positive utricle hair cells, with HCN1 and CNGA3 exhibiting the highest expression levels (Figure 5A). Importantly, expression profiling by qRT‒ PCR of single isolated utricle hair cells revealed the confined expression of CNGA3 in cluster I cells but not in cluster II cells in both *Gpr133*^-/-^ mice and their WT littermates (Figures 4B-4C, Figure 5B). Moreover, the direct association of GPR133 with CNGA3, but not with HCN1, HCN4, CNGA1 or CNGB1, was observed by coimmunoprecipitation in HEK293 cells or utricles, and the colocalization of GPR133 with CNGA3 was indicated by coimmunostaining (Figures 5C-5D, Figures S6A-S6C). Consistently, the application of the CNG inhibitor L-cis-diltiazem directly blocked the current change in GPR133-expressing cluster I utricle hair cells, but not in cluster II hair cells, in response to force stimulation via GPR133-M-beads (Figures 5E-5F)^56^. These results collectively indicate that coupling with CNGA3 underlies the modulation of membrane excitability of GPR133 in response to force sensation.

**Figure 5.**
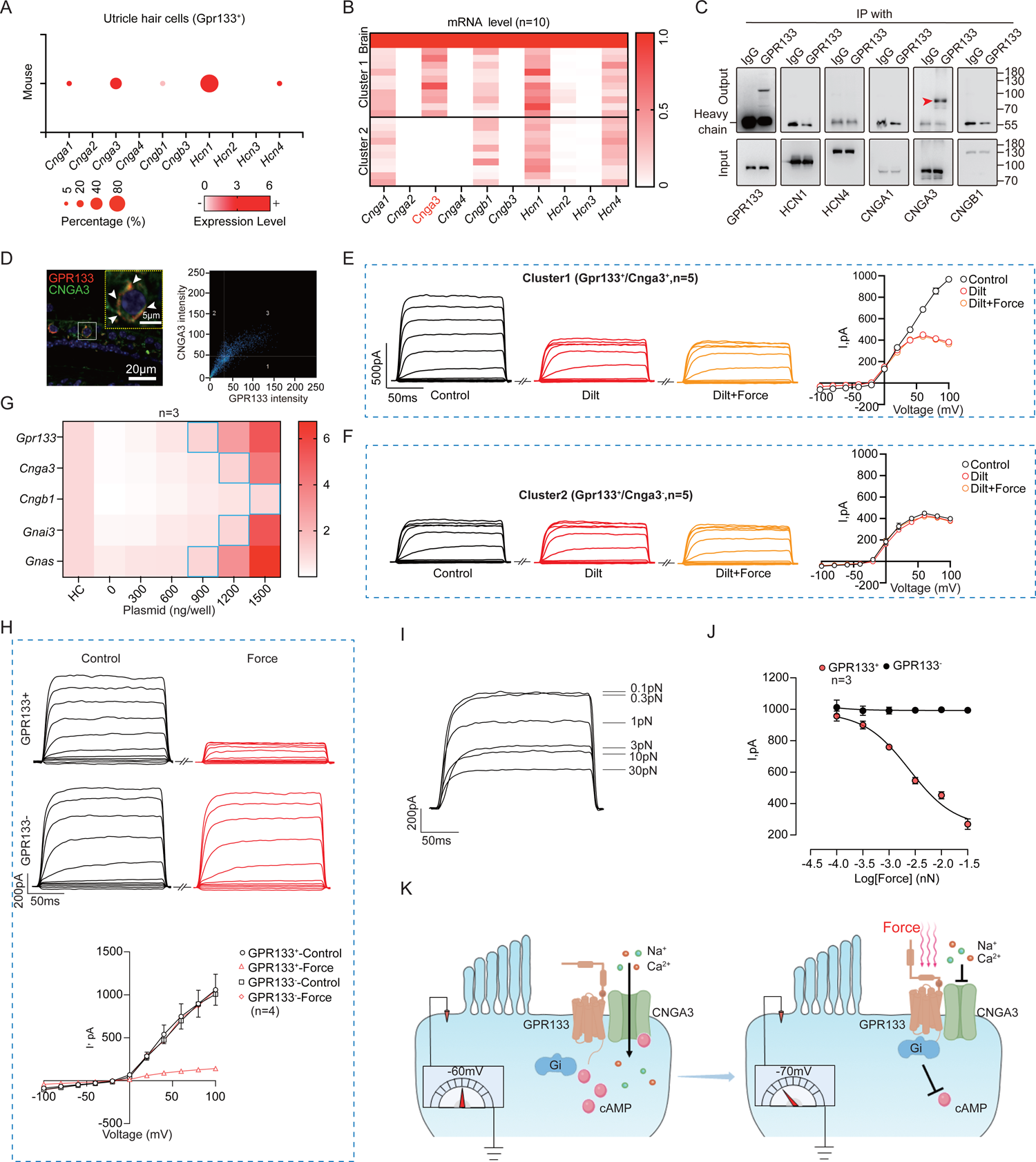
GPR133 regulates mechanoelectrical transduction through coupling to CNGA3. **(A)** Expression profiles of cyclic nucleotide–activated ion channels in GPR133-expressing utricle hair cells. Single cell RNA-seq datasets are from GSE71982. **(B)** mRNA levels of cyclic nucleotide–activated ion channels in *Gpr133* promoter-labeled cluster I and cluster II utricle hair cells examined by single cell qRT-PCR (n=10). **(C)** Co-immunoprecipitation of GPR133 with cyclic nucleotide–activated ion channels in the lysates of mouse utricle. Representative blotting from three independent experiments is shown. Data are correlated to Extended data Fig. 6b. **(D)** Left panel: Co-immunostaining of GPR133 (red) with CNGA3 (green) in utricle sections derived from WT mice (n = 3 mice per group). Scale bar: 20 μm and 5 μm for low low and high magnification view, respectively. Right panel: Pearson’s correlation analysis of GPR133 and CNGA3 fluorescence intensities. The Pearson’s correlation coefficient was 0.70. **(E-F)** Effect of the inhibitor of CNGA3-containing heteromeric channels L-cis-Diltiazem on the whole-cell current in cluster I (**E**) and cluster II (**F**) hair cells in response to force (3 pN) stimulation. **(G)** Screening the transfection amounts of plasmids encoding GPR133-CNG signaling components in HEK293 cells to enable similar expression levels of these components compared with those in native cluster I utricle hair cells (n=3). Data are correlated to Figure S6D-S6H. **(H)** Representative whole-cell currents (top panel) elicited by voltage steps between −100 mV and +100 mV and the corresponding I-V curves (bottom panel) in HEK293 cells overexpressing GPR133/CNGA3/CNGB1/Gi3/Gs or /CNGA3/CNGB1/Gi3/Gs in response to force (3 pN) stimulation (n=4). **(I-J)** Representative whole-cell currents (**I**) and the corresponding dose-response curves (**J**) in the heterologous system with or without GPR133 in response to force stimulation at a holding potential of +100 mV (n=3). **(K)** Schematic representation of the GPR133-CNGA3 functional coupling in mechanoelectrical transduction in utricle hair cells.

We next used in vitro HEK293 systems to reconstitute the signaling pathway of GPR133-regulated membrane excitability in response to force sensation. Notably, HEK293 cells had much lower expression levels of *Gpr133*, *Gnai3*, *Cngb1* and *Cnga3* than GPR133-expressing utricle hair cells (Figure 5G, Figures S6D-S6H). We therefore screened the plasmid amounts for transfection of HEK293 cells to ensure similar expression of these signaling components in HEK293 cells compared with that of native cluster I utricle hair cells (Figure 5G, Figures S6D-S6H). Application of force at 3 pN induced an approximately 66% reduction in cAMP levels in this in vitro reconstituted system pre-stimulated with forskolin, which was comparable with those recorded in utricle explants (Figures S6I). Consistently, an approximately 85% reduction in current amplitude at +100 mV in response to force stimulation via Flag-M-beads was readily detected in the above system but was abolished when GPR133 was absent (Figure 5H). At the holding potential of +100 mV, the force induced a dose-dependent decrease in current amplitude with an EC50 of 2.55±0.34 pN, similar to that recorded ex vivo (Figure 5I-5J). Therefore, these results supported a regulatory role of GPR133-CNGA3 functional coupling in mechanoelectrical transduction in utricle hair cells (Figure 5K).

### Specific balance-regulating function of GPR133 in vestibular hair cells

To further inspect the specific regulatory role of GPR133 in the vestibular system but not in other tissues, especially within the balance sensory circuit, we generated hair cell-specific *Gpr133*-knockout mice by crossing *Gpr133^fl/fl^*mice with inducible *Pou4f3-CreER^+/-^* transgenic mice because scRNA-seq data suggested restricted expression of GPR133 in *Pou4f3*^+^ utricle hair cells (Pou4f3 is the transcriptional target of ATOH1 and is commonly used as a marker of hair cells^57,58^) (Figures S7A-S7B). The ablation of GPR133 expression in the utricles of *Pou4f3-CreER^+/-^Gpr133^fl/fl^* mice was verified by both immunostaining and western blot analysis (Figure 6A, Figures S7C). In contrast, the expression of GPR133 in the vestibular nuclei within the brainstem was not affected in *Pou4f3-CreER^+/-^Gpr133^fl/fl^*mice, which was consistent with the lack of POU4F3 expression in this area (Figure 6B, Figure S7C).

**Figure 6.**
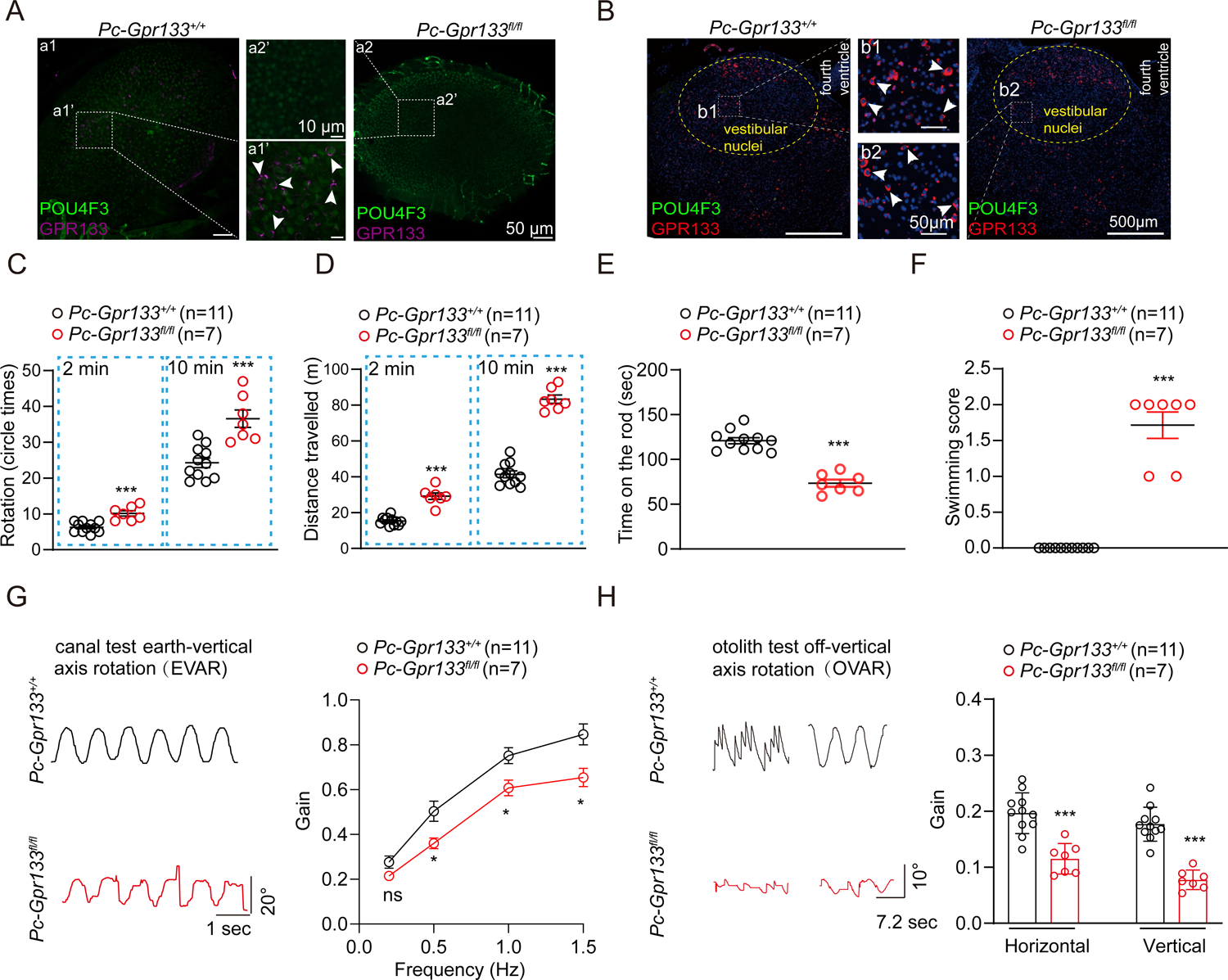
Specific ablation of GPR133 in hair cells impaired balance. **(A)** Immunostaining of GPR133 (red) and Pou4f3 (green) in utricle whole mounts derived from *Pou4f3-CreER^+/-^ Gpr133^fl/fl^* mice (referred to as *Pc-Gpr133 ^fl/fl^*) or *Pou4f3-CreER^+/-^Gpr133^+/+^*mice (referred to as *Pc-Gpr133^+/+^*). Enlarged images show the ablation of GPR133 in the hair cells of *Pc-Gpr133 ^fl/fl^* mice (n=3 mice per group). Scale bar: 50 μm and 10 μm for low and high magnification view, respectively. **(B)** Immunostaining of GPR133 (red) in the vestibular nuclei within the brainstem derived from *Pc-Gpr133 ^fl/fl^* mice or *Pc-Gpr133^+/+^* mice (n=3 mice per group). Scale bar: 500 μm and 50 μm for low and high magnification view, respectively. **(C-F)** Quantification of the circling activity (**C**), travelling activity (**D**), time on the rotating rod (**e**) and swimming scores (**f**) of *Pc-Gpr133 ^fl/fl^* mice and *Pc-Gpr133^+/+^* mice (n=11 mice per group). **(G)** Representative recording curves (left) and quantification of the VOR gain response of *Pc-Gpr133 ^fl/fl^* mice and *Pc-Gpr133^+/+^* mice to earth-vertical axis rotation (0.25-1.5 Hz, 40°/s peak velocity sinusoidal, whole-body passive rotation, n=11 mice per group). **(H)** Representative recording curves (left) and quantification of the VOR gain response of *Pc-Gpr133 ^fl/fl^* mice and *Pc-Gpr133^+/+^*mice to off-vertical axis rotation (50°/s, whole-body passive rotation, n=11 mice per group). (**C-H**) **P < 0.01; ***P < 0.001; ns, no significant difference. *Pc-Gpr133 ^fl/fl^* mice compared with *Pc-Gpr133^+/+^*mice. The bars indicate mean ± SEM values. All data were statistically analyzed using Student’s *t* test.

The *Pou4f3-CreER^+/-^Gpr133^fl/fl^* mice recapitulated the phenotypes of *Gpr133^-/-^* mice in most vestibular behavior studies, showing increased circling behaviors, decreased duration time on the rotarod and deteriorated swimming performance compared with those of the control *Pou4f3-CreER^+/-^Gpr133^+/+^* mice (Figures 6C-6F). Intriguingly, whereas the *Pou4f3-CreER^+/-^Gpr133^fl/fl^*mice showed more than 60% reduction in VOR gain in off-vertical axis rotation, which represents otolith functions, these mice displayed 20-30% reduction in VOR response to earth-vertical axis rotation at all tested frequencies, which reflects semicircular canal functions (Figures 6G-6H). These results suggested that GPR133 might play a more important role in the otolith organ. Because all (100%) of the hair cells in the utricles expressing GPR133 were *Pou4f3* positive and only 75% of hair cells in the semicircular canal were *Pou4f3* positive, we could not exclude the possibility that the residual GPR133 (25%) in the semicircular canal of *Pou4f3-CreER^+/-^Gpr133^fl/fl^*mice might account for less-affected canal-mediated VOR. An alternative explanation is that the equilibrioception mechanism in the semicircular canal is different from that in the otolith organs, which awaits further investigation.

### Potential mechanism of GPR133 activation by the force and structural basis of GPR133-Gi coupling

It has been proposed that the release of the *Stachel* sequence from the GAIN domain of aGPCRs is one possible mechanism mediating aGPCR activation in response to mechanical force stimulation^59^. Consistent with this hypothesis, only the GPR133 constructs with an intact GAIN domain could respond to force stimulation and regulate Gi activation and cAMP production, suggesting a critical role of the GAIN domain in force sensing (Figures 7A-7C, Figure S8A). To further delineate the dynamic process of force-induced GPR133 activation, we employed the monobromobimane (mBBr) labeling method combined with mass spectrometry (MS) analysis. mBBR specifically conjugates to the sulfur atom of cysteine^60^. We therefore mutated 6 residues within the *Stachel* sequence of GPR133 into cysteine and found that 3 mutants, including L544C, N546C and V554C, showed force-induced Gi3 activation comparable to that of the WT receptor when expressed at similar levels, thus indicating intact functionality in force sensation and Gi3 coupling (Figures 7D-7E, Figure S8B). These 3 mutants were purified, labeled with mBBr before and after force application, and analyzed by MS. Notably, a particular mutant, L544C, showed significantly increased mBBr labeling after force stimulation, suggesting force-induced exposure of the *Stachel* sequence from an intramolecular prebound state (Figure 7F). Consistent with this hypothesis, alanine mutations of selective *Stachel*-interacting residues in the GAIN domain, such as I483A or V508A/Y509A, led to increased basal Gi activity of the receptor but diminished force-sensing potential (Figures 7G-7I, Figure S8C), which likely mimicked an intermediate state during force-stimulated GPR133 activation (these residues were proposed by analyzing the full-length GPR133 structure by molecular dynamics simulation in our previous study^18^). Collectively, these data suggest that application of force to GPR133 induces exposure of the *Stachel* sequence within the GAIN domain, which increases the chance for binding of the *Stachel* sequence to the 7TM domain to facilitate receptor activation (Figure 7J).

**Figure 7.**
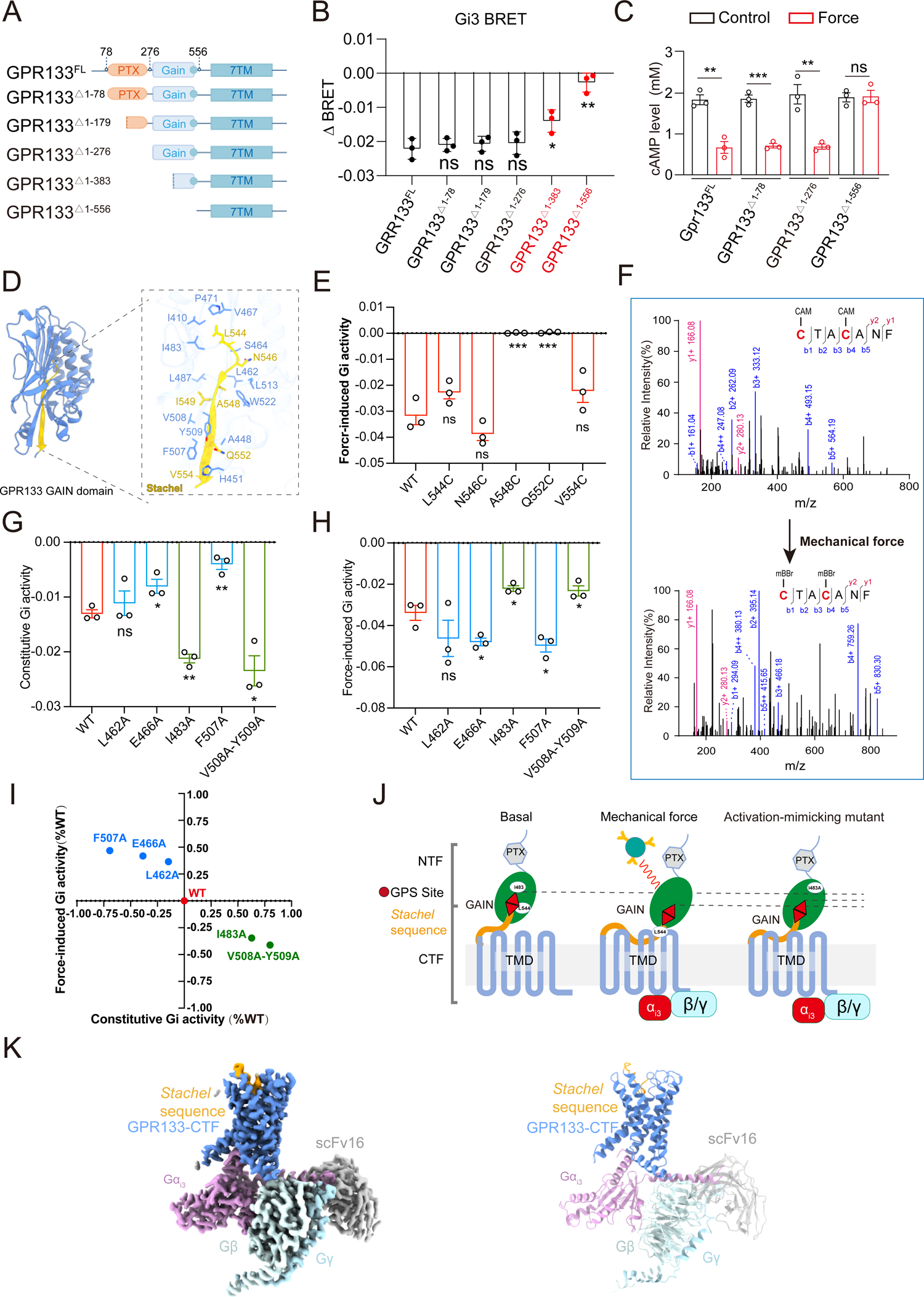
Structural basis for force-induced GPR133 activation. **(A)** Schematic representation of the generation of different N-terminal truncated constructs of GPR133. **(B)** Force (3 pN)-induced Gi3 activation in HEK293 cells transfected with WT full-length GPR133 or its N-terminal truncated mutants (n=3). **(C)** Force (3 pN)-induced cAMP reduction in HEK293 cells transfected with WT full-length GPR133 or its N-terminal truncated mutants (n=3). Transfection amounts of plasmids encoding GPR133-CNGA3 signaling components in HEK293 cells were adjusted to enable similar expression levels compared with those in native cluster I utricle hair cells. **(D)** Detailed interactions between the *Stachel* sequence and the GAIN domain according to simulated GPR133-FL structure by alphafold 2 according to our previous study. **(E)** Force (3 pN)-induced Gi3 activation in HEK293 cells transfected with WT full-length GPR133 or the cysteine mutations of *Stachel* residues (n=3). **(F)** Mass spectrum of the GPR133 *Stachel* sequence of GPR133-L544C before (top panel) and after (bottom panel) force stimulation. The carbamidomethylated cysteine residues were labelled with mBBr after force stimulation. **(G)** Constitutive Gi activity of WT GPR133 or GPR133 mutants located on the *Stachel*-GAIN domain interface. Constitutive Gi activity was calculated by subtracting the BRET value obtained in cells transfected only with Gi BRET probes from the BRET value obtained in cells transfected with GPR133 (WT or mutants) and Gi BRET probes (n=3). WT and the mutants of GPR133 were expressed at similar levels. Data are correlated to Figure S8C. **(H)** Force (3 pN) induced Gi activation through WT GPR133 or GPR133 mutants located on the *Stachel*-GAIN domain interface. Force-induced Gi activity was defined as the change in BRET values in cells transfected with GPR133 (WT or mutants) and Gi BRET probes in response to force stimulation (n=3). WT GPR133 and the mutants were expressed at similar levels. **(I)** Summary of the effects of mutations of *Stachel*-interacting GAIN residues on the basal and force-induced Gi activity measured by BRET assay. The X axis represents the increased constitutive Gi activity of GPR133 mutants using GPR133 WT as a reference (normalized to 1). A positive X value indicates the increased folds of constitutive Gi activity of indicated GPR133 mutant compared with WT GPR133. The Y axis represents the force-induced Gi activity of GPR133 mutants using that of GPR133 WT as a reference (normalized to 1). A positive Y value indicates increased force-induced Gi activity compared with WT GPR133. Mutants (I408A and V508A/Y509A) in the fourth quadrant displayed increased basal Gi activity but diminished force-sensing potential, thus mimicking an intermediate state during force-stimulated GPR133 activation (n=3). **(J)** Schematic representation of the potential mechanism underlying GPR133 activation after force sensing. GPR133 senses force stimulation induced by magnetic beads via its GAIN domain, which causes the release of the exposure of *Stachel* sequence from a pre-bound state, as evidenced by the increased mBBr labelling of L544C mutant. Consistently, alanine mutation of *Stachel*-interacting residues in GAIN domain, such as I483A, mimicked an intermediate state of force-stimulated GPR133 activation. **(K)** Cryo-EM density map and ribbon model of the GPR133-Gi3 complex (PDB ID: XXXX). GPR133, cornflower blue; *Stachel* sequence, orange; Gαi3, plum; Gβ, powder blue; Gγ, sky blue; scFv16, silver. (**B, E, G, H**) *P<0.05; **P<0.01; ns, no significant difference. GPR133 mutants compared with WT GPR133. (**C**) **P<0.01; ***P<0.001; ns, no significant difference. Force stimulated cells compared with control cells. The bars indicate mean ± SEM values. All data were statistically analyzed using one-way (B, E, G, H) or two-way (c) ANOVA with Dunnett’s post hoc test.

To provide a structural basis for the force-induced interaction between GPR133 and Gi and potential *Stachel* sequence-mediated Gi coupling, we solved the cryo-electron microscopy (cryo-EM) structure of the GPR133-CTF–Gi3-scFv16 complex with an overall resolution of 3.2 Å (Figure 7K, Figures S8D-S8F). The EM densities allowed assignment of most residues of the GPR133 7TM bundle and the heterotrimeric Gi3 protein (Figure 7K, Figure S8G). The overall structure of the GPR133 receptor bound by Gi3 is similar to that of the GPR133 receptor bound by Gs^18^, with a root-mean-square deviation (RMSD) of approximately 0.63 for the transmembrane domain (TMD). Similar to that in the GPR133-CTF–Gs complex structure, the *Stachel* sequence folded as a U shape, with the C-terminal β strand exposed to solution and the N-terminal α helix buried inside of the 7TM domain (Figure 8A). Whereas four residues of the hydrophobic interaction motif (HIM) of the *Stachel* sequence of GPR133 could be easily fit into the solved EM density, the side chain density for L550 (L^SS06^) of the HIM could not be clearly located, suggesting a potentially flexible conformation with less defined interactions (superscripts with numbers after “ss” (*Stachel* sequence) indicate the positions within the *Stachel* sequence) (Figures 8A-8F, Figures S9A-S9E). Moreover, the side chain of I549 (I^SS05^) of the HIM motif in the GPR133-CTF–Gi3 complex moved toward TM1 by approximately 2 Å compared with that in the GPR133-CTF–Gs complex, forming fewer interactions with N703 and W705 (Figure 8C, Figure S9F). In contrast, F547 (F^SS03^) of HIM rotated by approximately 30 degrees, forming more interactions with L550^ss06^, F643^3.40^ and the toggle switch W773^6.53^ (Figure 8D, Figure S9G). Consistent with these observations, mutations of *Stachel*-binding pocket residues, such as I713^5.36^, W714^5.37^, W773^6.53^, V777^6.57^, N795^7.46^ and F791^7.42^, significantly impaired force-induced Gi activation (Figure 8G, Figure S9H). In total, 3 residues of the HIM of the *Stachel* sequence impaired force-induced GPR133-Gi3 coupling (Figure 8H, Figure S9H). Notably, mutations of I549^SS05^A or L550^SS06^A significantly decreased force-induced Gs coupling to GPR133 but had no significant effects on force-induced Gi3 coupling. Conversely, the mutation of F547^SS03^A and L553^SS09^A of GPR133 decreased force-induced Gi3 activities more than Gs activities (Figure 8H).

**Figure 8.**
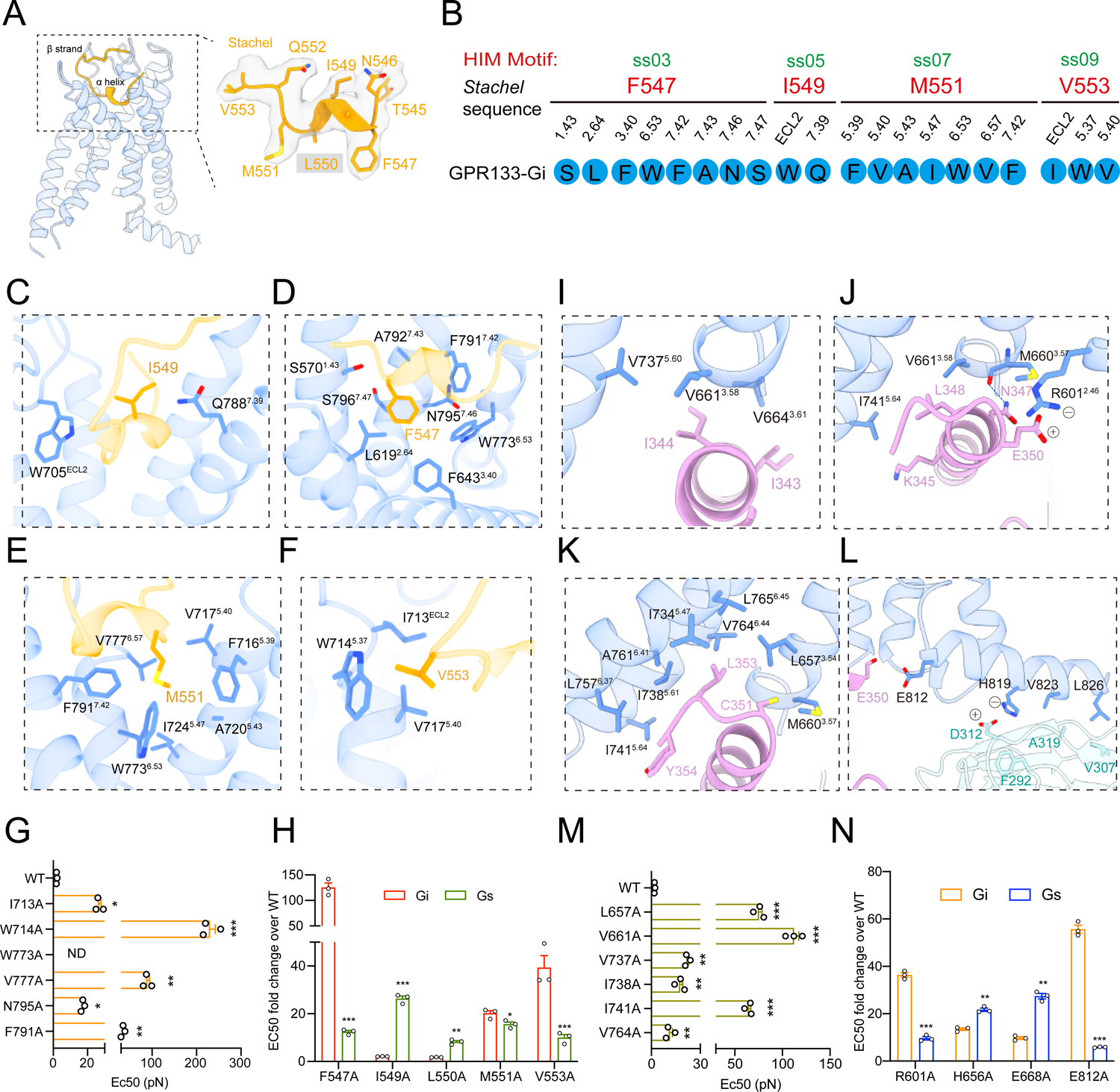
Potential structural basis of GPR133 activation and Gi coupling in response to force sensing. **(A)** Cryo-EM density map and the model of the *Stachel* sequence in GPR133-Gi3 complex. The side chain density for L550 was not clearly located, suggesting a potentially flexible conformation. **(B)** Binding pocket of the HIM motif residues of the GPR133 *Stachel* sequence. Residues in HIM motif are colored red and their interacting residues are circled in blue^18^. **(C-F)** Detailed interactions of the HIM residues, including I549 (**C**), F547 (**D**), M551 (**E**) and V553 (**F**), with the binding pocket of the GPR133-Gi3 complex. **(G)** Effects of mutations in GPR133 *Stachel* sequence binding pocket residues on the potency of force-induced Gi activity. WT GPR133 and the mutants were expressed at similar levels. Data are correlated to Figure S9h (n=3). **(H)** Effects of mutations in GPR133 *Stachel* sequence residues on the potency of force-induced Gi or Gs activity. The Y axis represents the fold change of the EC50 at force-induced Gs or Gi pathway of GPR133 mutants over that of WT GPR133. WT GPR133 and the mutants were expressed at similar levels. **(I-L)** Detailed interactions between GPR133 and heterotrimeric Gi3 protein. The interactions include hydrophobic interactions between receptor residues and I343/I344 in the α5 helix (**I**); GPR133 residues interacting with L345, N347and K348 in the α5 helix (**j**); GPR133 residues interacting with Y354 and L353 in α5 helix of Gαi (**K**); and extensive interactions between the helix 8 of GPR133 and Gα (E350) and Gβ (**L**). **(M)** Effects of mutations in G protein-contacting residues of GPR33 on the potency of force-induced Gi activity. WT GPR133 and the mutants were expressed at similar levels. Data are correlated to Figure S9O (n=3). **(N)** Effects of mutations in G protein-contacting residues of GPR33 on the potency of force-induced Gi or Gs activity. WT GPR133 and the mutants were expressed at similar levels. (**G, M**) *P<0.05; **P<0.01; ***P<0.001; ND, not detectable. GPR133 mutants compared with WT GPR133. (**H, N**) *P<0.05; **P<0.01; ***P<0.001. The potency of GPR133 mutants at Gs pathway compared with that at Gi pathway. The bars indicate mean ± SEM values. All data were statistically analyzed using one-way (G, M) or two-way (H, N) ANOVA with Dunnett’s post hoc test.

The structure of GPR133-CTF–Gi3 provided a potential Gi coupling mechanism of GPR133 activated by force application. The C-terminal α5 helix of Gαi3 inserted into the cavity created by the cytoplasmic ends of TM2-TM3 and TM5-TM7, similar to that of Gs in the GPR133-CTF–Gs complex structure (Figure 9I). I343^G.H5.15^, I344^G.H5.16^, K345^G.H5.17^, L348^G.H5.20^, L353^G.H5.25^ and Y354^G.H5.26^ of the α5 helix of Gαi3 constituted a long hydrophobic stretch and formed hydrophobic packings with L657^3.54^, V661^3.58^, V664^ICL2^, V734^5.57^, V737^5.60^, I738^5.61^, I741^5.64^, L757^6.37^, V764^6.44^ and L765^6.45^ of GPR133 (Figures 8I-8K). At the C-terminal H8 helix, E812^8.49^ formed polar interactions with the main chain carbonyl oxygen of E350^G.H5.22^ in the Gα α5 helix, while H819^8.56^, V823^8.60^ and L826^8.63^ of GPR133 formed charge‒charge or hydrophobic interactions with D312, F292, A319 and V307 of Gβ, respectively (Figure 8L). Despite sharing 16 common residues in GPR133 for contact with Gi3 compared with Gs, there were many fewer observed GPR133-Gi interface residues than GPR133-Gs interface residues (Figure S9J). For instance, the replacement of H41^G.S1.02^ of Gs by E33 in the structurally equivalent position of Gi3 resulted in loss of potential charge‒charge interactions with E668^ICL2^, and the replacement of Y391^G.H5.23^ of Gs with C351^G.H5.23^ of Gi3 resulted in loss of its hydrogen bond and hydrophobic packing with H656^3.53^(Figures S9K-S9L). In contrast, the replacement of Q390^G.H5.22^ of Gs with E350^G.H5.22^ of Gi3 enabled stronger interactions with R601 of GPR133 involved in salt bridge engagements (Figures S9M-S9N). Consistent with these observations, mutations of GPR133-Gi3 interface residues, such as L657^3.54^, V661^3.58^, V737^5.60^, I738^5.61^, I741^5.64^, and V764^6.44^, significantly impaired the force-induced Gi3 coupling to GPR133 (Figure 8M, Figure S9O). Mutations of E668A and H656A showed larger effects for Gs coupling than Gi3 coupling, whereas mutation of R601A had more effects for Gi3 coupling than Gs coupling (Figure 8N).

**Figure 9.**
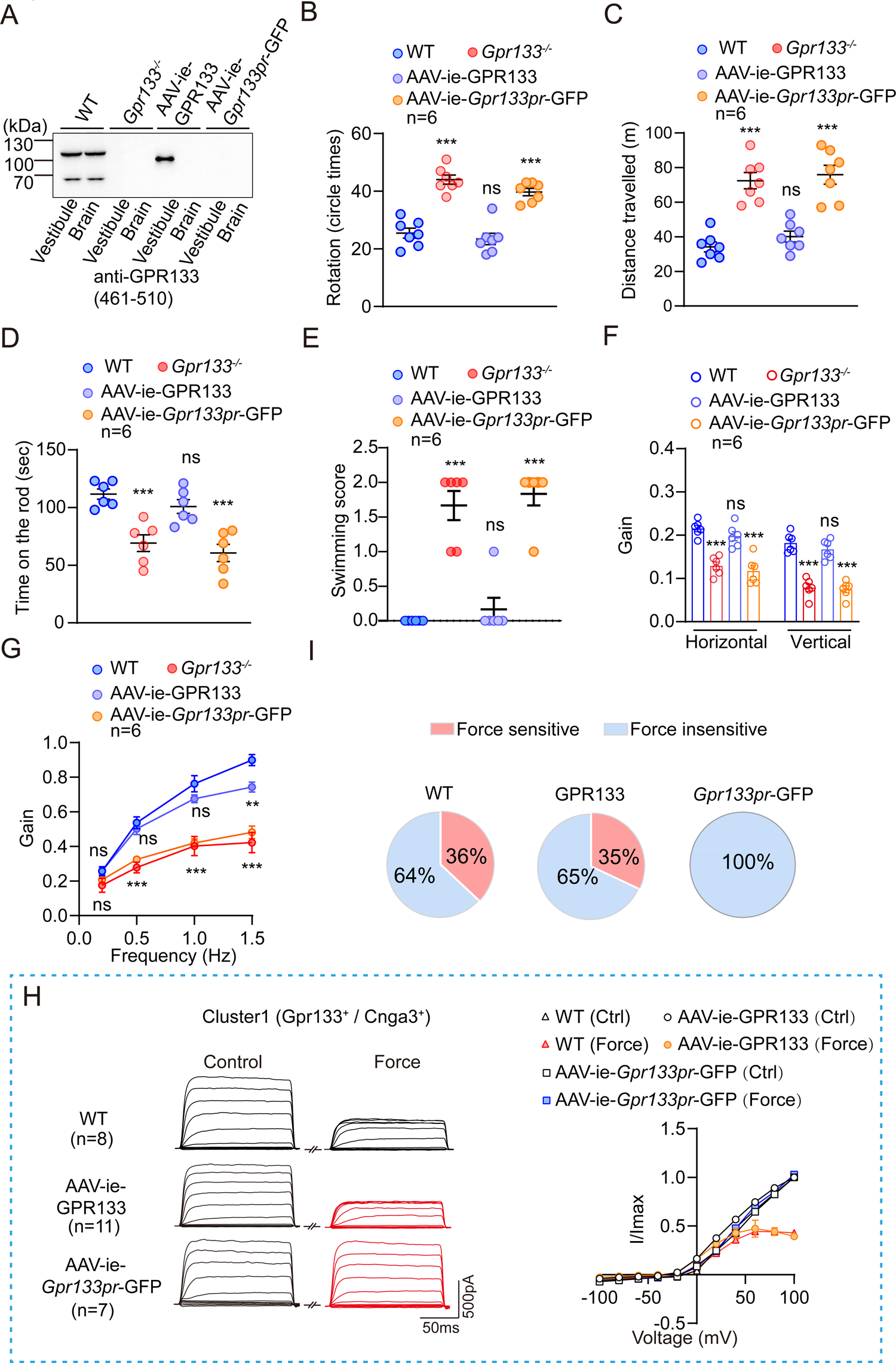
Specific expression of GPR133 rescues the vestibular function of *Gpr133^-/-^* mice. **(A)** Representative western blotting showing the expression of GPR133 in the membrane fractions of utricle or brainstem isolated from WT mice, *Gpr133^-/-^* mice, AAV-ie-GPR133-treated *Gpr133^-/-^* mice and AAV-ie-*Gpr133pr*-GFP-treated mice. The antibody against the N terminal (461-510 residues) of GPR133 was used, revealing the full-length (∼110 kDa) and a truncated GPR133 (∼90 kDa) in the WT and AAV-ie-GPR133-treated *Gpr133^-/-^* utricles, respectively. Representative blotting from three independent experiments were shown (n=3). **(B-C)** Quantification of the circling (B) and travelling activity (C) of WT mice, *Gpr133^-/-^* mice, AAV-ie-GPR133-treated *Gpr133^-/-^*mice and AAV-ie-*Gpr133pr*-GFP-treated mice in open-field test (n=6 mice per group). **(D)** Quantification of time on the rotating rod of WT mice, *Gpr133^-/-^* mice, AAV-ie-GPR133-treated *Gpr133^-/-^* mice and AAV-ie-*Gpr133pr*-GFP-treated mice in rotation test (n=6 mice per group). **(E)** Quantification of swimming scores of WT mice, *Gpr133^-/-^* mice, AAV-ie-GPR133-treated *Gpr133^-/-^* mice and AAV-ie-*Gpr133pr*-GFP-treated mice (n=6 mice per group). **(F-G)** Quantification of the VOR gain response of WT mice, *Gpr133^-/-^* mice, AAV-ie-GPR133-treated *Gpr133^-/-^*mice and AAV-ie-*Gpr133pr*-GFP-treated mice to off-vertical axis rotation (F) and earth-vertical axis rotation (G) (n=6 mice per group). **(H)** Representative whole-cell currents (left panel) elicited by voltage steps between −100 mV and +100 mV and the corresponding I-V curves (right panel) in cluster I utricle hair cells derived from WT mice, AAV-ie-GPR133-treated *Gpr133^-/-^*mice or AAV-ie-*Gpr133pr*-GFP-treated mice in response to force (3 pN) stimulation (n=8, 11, and 7 for WT, AAV-ie-GPR133, and AAV-ie-*Gpr133pr*-GFP group, respectively). **(I)** Quantification of force-sensitive and force-insensitive utricle hair cells derived from WT mice and *Gpr133^-/-^* mice injected with AAV-ie-GPR133 or AAV-ie-*Gpr133pr*-GFP. (**B-G**) ***P<0.001; ns, no significant difference. *Gpr133^-/-^* mice compared with WT mice. All data were statistically analyzed using one-way ANOVA with Dunnett’s post hoc test.

Taken together, these results suggest that release of the *Stachel* sequence from the *GAIN* domain and insertion of the *Stachel* sequence into the 7TM bundle of GPR133 mediates its force sensation process and that the cryo-EM structure provides preliminary knowledge of the force-induced GPR133-Gi3 coupling interface.

### Re-expression of GPR133 in the vestibule of *Gpr133^-/-^* mice rescues the equilibrioception

To further investigate the function of GPR133 in utricle hair cells, we performed in vivo rescue experiments. Due to the limited capacity of the AAV-ie (< 3.5 kb), we packaged the GPR133-GAIN-7TM sequences with a deletion of N-terminal 276 residues (with an intact GAIN domain, 7TM region and C-tail, referred to as AAV-ie-GPR133) instead of the full-length GPR133 coding sequence into the AAV-ie-*Gpr133pr*-GFP vector (GFP was fused to the C-terminus of GPR133-GAIN) (Figure 7A, Figure S10A). The total packaging size was approximately 3.2 kb (Gpr133 promoter 500 bp, Gpr133-GAIN 1.9 kb and GFP 720bp; both GPR133-GAIN and GFP expression was driven by the GPR133 promoter), which allowed successful AAV-ie production (Figure S10A). Notably, our structural and mutagenesis studies indicated that the GAIN domain is the main force-sensing part of GPR133 and the Gi-activating potential of GPR133-GAIN in response to force stimulation was comparable to that of the full-length receptor when expressed at the similar level, suggesting its functional integrity in force sensation (Figures 7B-7C). The AAV-ie-GPR133 was delivered into P3 *Gpr133^-/-^* mice through round window membrane (RWM) injection, with the AAV-ie-*Gpr133pr*-GFP as a negative control. The specific expression of GPR133 was observed in the utricle of *Gpr133^-/-^* mice injected with the AAV-ie-GPR133 but not in the brainstem at P40 as indicated by the western blotting and GFP fluorescence (Figure 9A, Figures S10A-S10C). Approximately 20% of the utricle hair cells of AAV-ie-GPR133-treated *Gpr133^-/-^* mice were labelled by

GFP, which is close to the proportion of GPR133-positive cells in WT mice revealed by GPR133 immunostaining (21%) or that in GPR133^GFP^ knock in mice (24.5%). Notably, the AAV-ie-GPR133-treated *Gpr133^-/-^* mice at P40, but not those treated by AAV-ie-*Gpr133pr*-GFP, showed significantly improved performance in the open field, rotarod and swimming tests, which were comparable to the levels of the WT mice (Figures 9B-9E). Moreover, the reduction in VOR gain of *Gpr133^-/-^* mice in off-vertical axis rotation test was fully reversed after AAV-ie-GPR133 injection (Figure 9F). Similarly, the VOR gains of AAV-ie-GPR133-treated *Gpr133^-/-^*mice were almost recovered to the levels of WT mice in earth-vertical axis rotation test except for the highest testing frequency of 1.5 Hz, at which the recovery efficiency was approximately 85% (Figure 9G). Electrophysiological analysis revealed that approximately 35% of GFP-labelled utricle hair cells in AAV-ie-GPR133-treated *Gpr133^-/-^*mice restored the force-induced excitability, as revealed by the significant current amplitude reduction in response to 3 pN force stimulation (Figures 9H-9I, Figure S10D-S10E). Both the proportion and current-voltage characteristics of the force-sensitive GPR133-expressiong utricle hair cells in AAV-ie-GPR133-treated *Gpr133^-/-^* mice resembled those observed for WT utricle cluster I hair cells (Figures 9H-9I). In contrast, none of the utricle hair cells in *Gpr133^-/-^* mice labelled by the AAV-ie-*Gpr133pr*-GFP showed detectable current amplitude alteration in response to force stimulation (Figures 9H-9I, Figure S10D). These data suggested that re-expression of GPR133 in the *Gpr133* promoter labeled utricle hair cells of *Gpr133^-/-^* mice could rectify the defect in force-induced excitability of the cells and rescue mostly impaired balance behaviors. Taken together, these results supported that the force-sensing GPR133 in the vestibular system specifically contributed to normal equilibrioception.

## Discussion

### GPR133 is essential for equilibrioception

Despite decades of intensive research, the mechanisms underlying equilibrioception and maintenance remain elusive. Membrane receptors, especially ion channels and GPCRs, are well known for their roles in basic sensing and maintenance for processes including but not limited to vision, olfaction, tasting, itching and touching^24,25,27,29,36^. Two types of ion channels, TMC1/2 and piezo2, were recently reported to be involved in the MET process in the utricle^10,12^. In particular, *Tmc1^-/-^Tmc2^-/-^* mice exhibited abnormal vestibular behaviors such as circling and head bobbing^10^. However, whether GPCRs also participate in equilibrioception is not known. Here, we provide direct evidence that a GPCR member, GPR133, expressed at the surface of vestibular hair cells, is essential for equilibrioception. Both heterozygous *Gpr133*^+/-^ mice (at P40) and homozygous *Gpr133*^-/-^ mice (up to P150) showed normal anatomic features and hair bundle polarity in the utricle maculae but profound vestibular behavior dysfunctions and impaired vestibulo-ocular reflexes. In contrast, the hearing function of *Gpr133*-deficient mice was not affected, as evidenced by the normal hearing threshold in the ABR and DPOAE measurements. These results suggested that we identified GPR133 as a specific GPCR member governing balance but not hearing function. Both *Gpr133*^+/-^ and *Gpr133*^-/-^ mice showed impaired compensatory eye movement in the VOR test. The VOR eye movement response reflects a combination of input, central processing and output of the balance signal. Importantly, whereas conditional ablation of GPR133 in hair cells recapitulated the deteriorated balance behaviors of the global *Gpr133*-deficient mice, specific re-expression of GPR133 in vestibular hair cells of *Gpr133*^-/-^ mice rescued the equilibrioception, thus suggesting a specific role of GPR133 in regulating the balance signal input in the vestibular system. To understand the exact role of GPR133 in the vestibular system and equilibrioception, direct measurements of peripheral vestibular functions, such as vestibular evoked potential (VsEP), by *Gpr133* conditional knockout mice are required in future studies.

### GPR133 is a sensor for mechano-second messenger transduction (MSMT) in the utricle

By leveraging a magnetic tweezer system connected with the GPCR biosensor platform, we were able to observe that mechanical force activated both Gs and Gi signaling pathways downstream of GPR133. In tissues, force stimulation significantly decreased the basal second messenger cAMP levels in vestibular hair cells by activating GPR133, indicating a mechano-second-messenger-transduction (MSMT) process in utricles. These results suggest that balance/force sensing in utricles may convert to changes in intracellular second messengers to modulate physiological processes. Notably, it has been reported that olfactory receptors (ORs) direct the targeting of olfactory sensory neuron axons to the olfactory bulb by regulating the Gs-cAMP signal, which controls the transcription of genes encoding axon guidance molecules and guides glomerular positioning^61^. In auditory and vestibular systems, the Gi protein is critical for stereocilia development and the establishment of hair cell polarity^37,62–64^. Therefore, it is slightly surprising that no significant morphological or anatomical abnormalities were observed for the utricle from *Gpr133^-/-^* mice at least at P40 and *Gpr133^+/-^* mice even at P150. We speculated that the short-term Gi signal defect in *Gpr133-*deficient mice might be compensated by other Gi-coupled receptors. In addition to GPR133, it is noteworthy that four other adhesion GPCRs expressed in the vestibular system, including GPR126, LPHN2, LPHN3 and VLGR1, were able to activate Gi signaling after force sensation. Although the knockout phenotypes indicated that VLGR1 is dispensable for balance, the regulatory roles of GPR126, LPHN2 and LPHN3 require further inspection. It would be interesting to investigate the synergetic effects of this group of MSMT-regulating GPCRs on vestibular system development.

### GPR133 mediates MET through coupling to CNGA3

In particular, we have revealed that GPR133 in vestibular hair cells functionally couples to CNGA3 to convert force sensation to changes in membrane excitability. Therefore, we have identified a particular GPCR, instead of the well-known MET sensor ion channels, that can mediate the MET process in the utricle. Interestingly, the present results let us recall the light-electrical transduction (LET) in photoreceptor cells and the odorant-electrical transduction (OET) in the olfactory epithelium, which were also mediated by coupling of separate GPCR groups to distinct CNG channels^65–67^.

Notably, kinetics analysis of GPR133-CNGA3 coupling and activation revealed that the delayed time between force stimulation and the change in hair cell membrane potential was 658.0 ± 11.14 ms, which is significantly longer than the time recorded in mechanically activated ion channels, such as piezo2 and TACAN (usually with millisecond latency)^68,69^. This slow response kinetics might be due to the diffusion limit of cAMP in response to force stimulation. Therefore, the equilibrioception regulated by GPR133 seems to be a relatively long process that is different from other instantaneous sensing processes, such as touch and pain, which have been demonstrated to be regulated by mechanically activated ion channels. Although we observed that Gi activation downstream of force sensation by GPR133 caused hyperpolarization of the hair cells, we could not exclude that Gs activation downstream of GPR133 in certain hair cells may cause depolarization after force sensation by GPR133 because more Gs expression over Gi expression may be found in specific utricle hair cells and GPR133-G protein subtype coupling is exquisitely regulated by differential expression of G protein subtype levels. For example, force-induced GPR133 activation led to an increase in intracellular cAMP in HEK293 model cells^18^. Further high-quality single-cell transcriptome analysis and patch-clamp recordings with more hair cells expressing GPR133 in utricles are necessary to evaluate this potential.

### Potential mechanism connecting the MET signal of GPR133 to equilibrioception

Although we could not provide direct evidence that the changes in the membrane excitability of utricle hair cells produced by force sensing of GPR133 could be transmitted to vestibular nuclei in the pons and medulla for equilibrioception, there remains a considerable possibility that this could happen. Multiple lines of evidence have suggested that TMC1 and TMC2 are candidate pore-forming subunits of the MET channel located at the tip of stereocilia responsible for both hearing and balance^11,70^. This raises the question of why we need another mechanosensor in utricle hair cells to convert the mechanical force to changes in membrane excitability. In one possible model, the combination of force sensation by TMC1/TMC2 and GPR133 provides complete temporal and positional information. Notably, the force sensation by TMC1/2 and GPR133 have different temporal windows and expression patterns. By examining recent scRNA-seq data, we noticed that GPR133 was expressed in more than one-third of TMC1- or TMC2-positive vestibular hair cells, which was significantly higher than the proportion of GPR133-expressing cells among hair cells (approximately 20%)^71^. In contrast, approximately 50% of hair cells expressed TMC1/TMC2 but showed no GPR133 expression. Therefore, hair cells in utricles could be divided into at least three types according to TMC1/2 and GPR133 expression: TMC1/2^+^GPR133^-^ (type A) cells (∼50%), TMC1/2^+^GPR133^+^ (type B) cells (∼20%) and TMC1/2^-^GPR133^+^ (type C) cells (∼1%) (Figure S10G). We speculated that GPR133 might function as a ‘modem’ located in type B cells, modulating TMC-mediated electrical signals and transmitting ‘processed data’ into the central nervous system through afferent neurons. This MET information generated by “type B” hair cells might be compared with “type A” hair cells reported to the central nervous system to acquire more precise spatial and temporal information, especially considering that the time frame for GPR133-mediated MET is significantly slower than those generated by TMC1/2. In addition, the electrical signal generated by GPR133 in “type C” hair cells might directly connect to the central nervous system.

The details of these connections may involve the release of neurotransmitters from these cells and production of electrical signals in specific groups of afferent neurons, which may be studied through a combination of lineage-tracing technology and spatial multiomics. It will be of particular interest to investigate how GPR133-regulated electrical signals are transmitted from vestibular hair cells to the brain and how the brain modulates the hardwired circuit to control balance behaviors.

When examining the expression pattern of GPR133 in utricle hair cells, we found that GPR133 expression was also enriched at the basal membrane, where hair cells might contact supporting cells or afferent neurons. Therefore, it is also possible that GPR133 interacts with certain membrane receptors to bridge neighboring cells. Head motion might induce relative movement between these cells, which triggers *cis* signaling in hair cells by mechanically manipulating GPR133 activity. This speculation recalls the outer hair cells (OHCs) in the mammalian cochlea, which convert membrane potential changes into somatic longitudinal motility and amplify sound-induced electrical signals^72,73^. Therefore, it will also be of interest to investigate whether a similar mechanism exists in the vestibular system that potentially regulates the shape or motility of some GPR133-expressing hair cells and fine-tunes the sensitivity of equilibrioception.

### Structural basis of the force sensation by GPR133 and subsequent Gi3 coupling

In addition to sensing force through ion channels, sensing force by GPCRs participates in many important physiological processes. Here, our chemical labeling and MS data suggested that force application induced dislodgement of the *Stachel* sequence from the GAIN domain. Further cryo-EM structures and mutagenesis analysis not only provided details of the GPR133-Gi3 interface but also indicated that three residues of the HIM of the *Stachel* sequence mediated force-induced Gi coupling to GPR133. In particular, two of them, F547^SS03^ and L553^SS09^, contributed more to force-induced Gi3 activation than to force-induced Gs activation. Therefore, GPR133-F547^SS03A^ mutant knock-in mice may be useful to specify the exact role of GPR133-Gi signaling in mechanical force sensation and balance.

### Limitations of the current study and future perspectives

Notably, the essential role of GPR133 in equilibrioception may also be dependent on the expression of GPR133 in the extravestibular system, such as in the vestibular nuclei within the brainstem. The definition of the specific markers for the vestibular nuclei in future studies will be helpful to investigate the functional role of GPR133 in the sensory circuit. In addition, the specific role of CNGA3, which is required in MET signal generated by GPR133 in force sensation, in equilibrioception is not defined in our current study. Future studies using conditional knockout mice in which CNGA3 is specifically ablated in vestibular hair cells (e.g., by crossing *CNGA3 ^fl/fl^*mice with *Pou4f3-Cre* mice) could be exploited to explore the role of CNGA3.

Despite the high prevalence of vertigo and dizziness, which are among the most common complaints in neurological patients, selective clinical therapies for balance dysfunction are still lacking. The identification of the essential role of GPR133 in equilibrioception in the current study may provide an important therapeutic avenue for the treatment of vertigo and dizziness because GPCR ligands account for approximately 35% of current clinical drugs. In vitro and in silico screening and rational design of GPR133-selective ligands based on the structure of the GPR133-Gi complex acquired here may lead to useful therapeutic strategies.

## Data availability

All data are included in the main text or the extended materials. The cryo-EM density maps and atomic coordinates of GPR133-Gi3 complex have been deposited at the Electron Microscopy Data Bank (EMDB) and Protein Data Bank (PDB) under accession codes EMD-**** and ****, respectively. All other data are available when requested from the corresponding authors.

## Acknowledgments

The cryo-EM data were collected at the Cryo-Electron Microscopy Research Center at South University of Science and Technology of China. This work was supported by National Key R&D Program of China (2019YFA0904200 to J.-P.S. and P.X., 2021YFA1101300 to R.-J.C.), National Science Fund for Distinguished Young Scholars Grant (81825022 to J.-P.S., 82225011 to X.Y.), National Science Fund for Excellent Young Scholars (32222038 to P.X.), National Natural Science Foundation of China (32130055 and 81773704 to J.-P.S., 92057121 to X.Y., 82030029 to R.-J.C., 82271175 to X.-L.F.), the Key Research Project of the Beijing Natural Science Foundation, China (Z200019 to J.-P.S.), Major Fundamental Research Program of Natural Science Foundation of Shandong Province, China (ZR2020ZD39 to J.-P.S., ZR2021ZD18 to X.Y.), Shandong Provincial Natural Science Fund for Excellent Young Scholars (ZR2021YQ18 to P.X.)

## Author contributions

J.-P.S., X.Y. and R.-J.C initiated, designed and supervised the overall project. J.-P.S., X.Y. and Z.Y. starting the screening of mechanosensitive GPCRs in vestibular and cochlear systems from 2019. Z.Y. and J.-P.S. developed the magnetic force stimulation assay. X.Y. designed and supervised all electrophysiology experiments. Z.Y., S.-H.Z. and M.-W.W. performed the magnetic force stimulation assay. S.-H.Z., Z.Y., Q.-Y.Z. and H.L. performed the electrophysiological experiments on utricle hair cells and in the HEK293 reconstitution system supervised by J.-P.S. and X.Y.. Z.Y., S.-H.Z., M.-W.W., X.-L.F., Q.-Y.Z. and Z.-C.S. performed vestibular and auditory behavior studies. Z.Y., S.-H.Z., M.-W.W., X.-L.F., Q.-Y.Z. and Z.-C.S. isolated mice utricle and performed immunofluorescence studies and ex vivo experiments. Z.Y., S.-H.Z., M.-W.W. and X.-L.F. performed SEM experiment. Z.Y., M.-W.W., S.-H.Z., Q.-Y.Z., and Y.Z. analyzed the scRNA sequencing data supervised by J.-P.S.. Z.Y., M.-W.W. and S.-H.Z. performed coimmunoprecipitation and western blotting analyses. J.-P.S. and P.X. designed and guided all structural analyses. Y.-Q.P., R.-J.Z., and P.X. purified GPR133-Gi complex, prepared the samples for cyro-EM. Y.-Q.P., P.X. and R.-J.Z. collected the cryo-EM data and performed cryo-EM map calculation, model building and refinement. J.-P.S. and P.X. designed all the GPR133 mutants, and Y.-Q.P., R.-J.Z. and Y.-T.X. generated the constructs and performed the cell-based assays. Y.L. performed mBBr labeling and mass spectrometry. W.Q. provided the Gauss/Tesla meter for the quantification of magnetic force. F.Y. provided important insights into the animal experiment design and result interpretation. J.-P.S., X.Y., R.-J.C., F.Y., Z.Y., S.-H.Z. and M.-W.W. participated in data analysis and interpretation. Z.Y., S.-H.Z., M.-W.W., Q.-Y.Z., Z.-C.S., B.-W.B., and X.C. prepared the figures. Z.Y., J.-P.S., X.Y., R.-J.C and F.Y. wrote the manuscript. All the authors have seen and commented on the manuscript.

## Submission history

The manuscript was submitted on Sep 20, 2023 and was sent for peer review on Sep 26, 2023 by the journal editor.

## References

1 Cullen, K. E. Vestibular processing during natural self-motion: implications for perception and action. Nat Rev Neurosci 20, 346–363, doi:10.1038/s41583-019-0153-1 (2019).

2 Brandt, T. & Dieterich, M. The dizzy patient: don’t forget disorders of the central vestibular system. Nat Rev Neurol 13, 352–362, doi:10.1038/nrneurol.2017.58 (2017).

3 Zheng, W. & Holt, J. R. The Mechanosensory Transduction Machinery in Inner Ear Hair Cells. Annu Rev Biophys 50, 31–51, doi:10.1146/annurev-biophys-062420-081842 (2021).

4 Pan, B. et al. Gene therapy restores auditory and vestibular function in a mouse model of Usher syndrome type 1c. Nat Biotechnol 35, 264–272, doi:10.1038/nbt.3801 (2017).

5 Ono, K. et al. Retinoic acid degradation shapes zonal development of vestibular organs and sensitivity to transient linear accelerations. Nat Commun 11, 63, doi:10.1038/s41467-019-13710-4 (2020).

6 Everett, L. A., Morsli, H., Wu, D. K. & Green, E. D. Expression pattern of the mouse ortholog of the Pendred’s syndrome gene (Pds) suggests a key role for pendrin in the inner ear. Proc Natl Acad Sci U S A 96, 9727–9732, doi:10.1073/pnas.96.17.9727 (1999).

7 Kurima, K. et al. TMC1 and TMC2 Localize at the Site of Mechanotransduction in Mammalian Inner Ear Hair Cell Stereocilia. Cell Rep 12, 1606–1617, doi:10.1016/j.celrep.2015.07.058 (2015).

8 Xiong, W. et al. TMHS is an integral component of the mechanotransduction machinery of cochlear hair cells. Cell 151, 1283–1295, doi:10.1016/j.cell.2012.10.041 (2012).

9 Zhao, B. et al. TMIE is an essential component of the mechanotransduction machinery of cochlear hair cells. Neuron 84, 954–967, doi:10.1016/j.neuron.2014.10.041 (2014).

10 Kawashima, Y. et al. Mechanotransduction in mouse inner ear hair cells requires transmembrane channel-like genes. J Clin Invest 121, 4796–4809, doi:10.1172/JCI60405 (2011).

11 Jeong, H. et al. Structures of the TMC-1 complex illuminate mechanosensory transduction. Nature 610, 796–803, doi:10.1038/s41586-022-05314-8 (2022).

12 Wu, Z. et al. Mechanosensory hair cells express two molecularly distinct mechanotransduction channels. Nat Neurosci 20, 24–33, doi:10.1038/nn.4449 (2017).

13 Scimia, M. C. et al. APJ acts as a dual receptor in cardiac hypertrophy. Nature 488, 394–398, doi:10.1038/nature11263 (2012).

14 Zou, Y. et al. Mechanical stress activates angiotensin II type 1 receptor without the involvement of angiotensin II. Nat Cell Biol 6, 499–506, doi:10.1038/ncb1137 (2004).

15 Tang, W., Strachan, R. T., Lefkowitz, R. J. & Rockman, H. A. Allosteric modulation of beta-arrestin-biased angiotensin II type 1 receptor signaling by membrane stretch. J Biol Chem 289, 28271–28283, doi:10.1074/jbc.M114.585067 (2014).

16 Chachisvilis, M., Zhang, Y. L. & Frangos, J. A. G protein-coupled receptors sense fluid shear stress in endothelial cells. Proc Natl Acad Sci U S A 103, 15463–15468, doi:10.1073/pnas.0607224103 (2006).

17 Yeung, J. et al. GPR56/ADGRG1 is a platelet collagen-responsive GPCR and hemostatic sensor of shear force. Proc Natl Acad Sci U S A 117, 28275–28286, doi:10.1073/pnas.2008921117 (2020).

18 Ping, Y. Q. et al. Structural basis for the tethered peptide activation of adhesion GPCRs. Nature 604, 763–770, doi:10.1038/s41586-022-04619-y (2022).

19 Mitgau, J. et al. The N Terminus of Adhesion G Protein-Coupled Receptor GPR126/ADGRG6 as Allosteric Force Integrator. Front Cell Dev Biol 10, 873278, doi:10.3389/fcell.2022.873278 (2022).

20 Xiao, P. et al. Tethered peptide activation mechanism of the adhesion GPCRs ADGRG2 and ADGRG4. Nature 604, 771–778, doi:10.1038/s41586-022-04590-8 (2022).

21 Liu, D. et al. CD97 promotes spleen dendritic cell homeostasis through the mechanosensing of red blood cells. Science 375, eabi5965, doi:10.1126/science.abi5965 (2022).

22 Bassilana, F., Nash, M. & Ludwig, M. G. Adhesion G protein-coupled receptors: opportunities for drug discovery. Nat Rev Drug Discov 18, 869–884, doi:10.1038/s41573-019-0039-y (2019).

23 Bautista, D. M. et al. The menthol receptor TRPM8 is the principal detector of environmental cold. Nature 448, 204–208, doi:10.1038/nature05910 (2007).

24 Coste, B. et al. Piezo1 and Piezo2 are essential components of distinct mechanically activated cation channels. Science 330, 55–60, doi:10.1126/science.1193270 (2010).

25 Fesenko, E. E., Kolesnikov, S. S. & Lyubarsky, A. L. Induction by cyclic GMP of cationic conductance in plasma membrane of retinal rod outer segment. Nature 313, 310–313, doi:10.1038/313310a0 (1985).

26 Stryer, L. Cyclic GMP cascade of vision. Annu Rev Neurosci 9, 87–119, doi:10.1146/annurev.ne.09.030186.000511 (1986).

27 Buck, L. & Axel, R. A novel multigene family may encode odorant receptors: a molecular basis for odor recognition. Cell 65, 175–187, doi:10.1016/0092-8674(91)90418-x (1991).

28 Kaupp, U. B. & Seifert, R. Cyclic nucleotide-gated ion channels. Physiol Rev 82, 769–824, doi:10.1152/physrev.00008.2002 (2002).

29 Zhao, G. Q. et al. The receptors for mammalian sweet and umami taste. Cell 115, 255–266, doi:10.1016/s0092-8674(03)00844-4 (2003).

30 Tu, Y. H. et al. An evolutionarily conserved gene family encodes proton-selective ion channels. Science 359, 1047–1050, doi:10.1126/science.aao3264 (2018).

31 Scholz, N. et al. The adhesion GPCR latrophilin/CIRL shapes mechanosensation. Cell Rep 11, 866–874, doi:10.1016/j.celrep.2015.04.008 (2015).

32 Petersen, S. C. et al. The adhesion GPCR GPR126 has distinct, domain-dependent functions in Schwann cell development mediated by interaction with laminin-211. Neuron 85, 755–769, doi:10.1016/j.neuron.2014.12.057 (2015).

33 Boyden, S. E. et al. Vibratory Urticaria Associated with a Missense Variant in ADGRE2. N Engl J Med 374, 656–663, doi:10.1056/NEJMoa1500611 (2016).

34 Alper, S. L. & Sharma, A. K. The SLC26 gene family of anion transporters and channels. Mol Aspects Med 34, 494–515, doi:10.1016/j.mam.2012.07.009 (2013).

35 Cheng, J. et al. Autonomous sensing of the insulin peptide by an olfactory G protein-coupled receptor modulates glucose metabolism. Cell Metab 34, 240–255 e210, doi:10.1016/j.cmet.2021.12.022 (2022).

36 Yang, F. et al. Structure, function and pharmacology of human itch receptor complexes. Nature 600, 164–169, doi:10.1038/s41586-021-04077-y (2021).

37 Mauriac, S. A. et al. Defective Gpsm2/Galpha(i3) signalling disrupts stereocilia development and growth cone actin dynamics in Chudley-McCullough syndrome. Nat Commun 8, 14907, doi:10.1038/ncomms14907 (2017).

38 Jiang, T., Kindt, K. & Wu, D. K. Transcription factor Emx2 controls stereociliary bundle orientation of sensory hair cells. Elife 6, doi:10.7554/eLife.23661 (2017).

39 Mecca, A. A., Caprara, G. A. & Peng, A. W. cAMP and voltage modulate rat auditory mechanotransduction by decreasing the stiffness of gating springs. Proc Natl Acad Sci U S A 119, e2107567119, doi:10.1073/pnas.2107567119 (2022).

40 McGee, J. et al. The very large G-protein-coupled receptor VLGR1: a component of the ankle link complex required for the normal development of auditory hair bundles. J Neurosci 26, 6543–6553, doi:10.1523/JNEUROSCI.0693-06.2006 (2006).

41 Hu, Q. X. et al. Constitutive Galphai coupling activity of very large G protein-coupled receptor 1 (VLGR1) and its regulation by PDZD7 protein. J Biol Chem 289, 24215–24225, doi:10.1074/jbc.M114.549816 (2014).

42 Yang, H., Xie, X., Deng, M., Chen, X. & Gan, L. Generation and characterization of Atoh1-Cre knock-in mouse line. Genesis 48, 407–413, doi:10.1002/dvg.20633 (2010).

43 Bermingham, N. A. et al. Math1: an essential gene for the generation of inner ear hair cells. Science 284, 1837–1841, doi:10.1126/science.284.5421.1837 (1999).

44 Hardisty-Hughes, R. E., Parker, A. & Brown, S. D. A hearing and vestibular phenotyping pipeline to identify mouse mutants with hearing impairment. Nat Protoc 5, 177–190, doi:10.1038/nprot.2009.204 (2010).

45 Li, A., Xue, J. & Peterson, E. H. Architecture of the mouse utricle: macular organization and hair bundle heights. J Neurophysiol 99, 718–733, doi:10.1152/jn.00831.2007 (2008).

46 Eatock, R. A. & Songer, J. E. Vestibular hair cells and afferents: two channels for head motion signals. Annu Rev Neurosci 34, 501–534, doi:10.1146/annurev-neuro-061010-113710 (2011).

47 Ji, Y. R. et al. Function of bidirectional sensitivity in the otolith organs established by transcription factor Emx2. Nat Commun 13, 6330, doi:10.1038/s41467-022-33819-3 (2022).

48 Folts, C. J., Giera, S., Li, T. & Piao, X. Adhesion G Protein-Coupled Receptors as Drug Targets for Neurological Diseases. Trends Pharmacol Sci 40, 278–293, doi:10.1016/j.tips.2019.02.003 (2019).

49 Zhang, D. L. et al. Gq activity- and beta-arrestin-1 scaffolding-mediated ADGRG2/CFTR coupling are required for male fertility. Elife 7, doi:10.7554/eLife.33432 (2018).

50 An, W. et al. Progesterone activates GPR126 to promote breast cancer development via the Gi pathway. Proc Natl Acad Sci U S A 119, e2117004119, doi:10.1073/pnas.2117004119 (2022).

51 Luo, R. et al. G protein-coupled receptor 56 and collagen III, a receptor-ligand pair, regulates cortical development and lamination. Proc Natl Acad Sci U S A 108, 12925–12930, doi:10.1073/pnas.1104821108 (2011).

52 Paavola, K. J., Stephenson, J. R., Ritter, S. L., Alter, S. P. & Hall, R. A. The N terminus of the adhesion G protein-coupled receptor GPR56 controls receptor signaling activity. J Biol Chem 286, 28914–28921, doi:10.1074/jbc.M111.247973 (2011).

53 Kishore, A. & Hall, R. A. Disease-associated extracellular loop mutations in the adhesion G protein-coupled receptor G1 (ADGRG1; GPR56) differentially regulate downstream signaling. J Biol Chem 292, 9711–9720, doi:10.1074/jbc.M117.780551 (2017).

54 Sun, Y. et al. Optimization of a peptide ligand for the adhesion GPCR ADGRG2 provides a potent tool to explore receptor biology. J Biol Chem 296, 100174, doi:10.1074/jbc.RA120.014726 (2021).

55 Tan, F. et al. AAV-ie enables safe and efficient gene transfer to inner ear cells. Nat Commun 10, 3733, doi:10.1038/s41467-019-11687-8 (2019).

56 Feketa, V. V., Nikolaev, Y. A., Merriman, D. K., Bagriantsev, S. N. & Gracheva, E. O. CNGA3 acts as a cold sensor in hypothalamic neurons. Elife 9, doi:10.7554/eLife.55370 (2020).

57 Vahava, O. et al. Mutation in transcription factor POU4F3 associated with inherited progressive hearing loss in humans. Science 279, 1950–1954, doi:10.1126/science.279.5358.1950 (1998).

58 Yu, H. V. et al. POU4F3 pioneer activity enables ATOH1 to drive diverse mechanoreceptor differentiation through a feed-forward epigenetic mechanism. Proc Natl Acad Sci U S A 118, doi:10.1073/pnas.2105137118 (2021).

59 Wilde, C., Mitgau, J., Suchy, T., Schoneberg, T. & Liebscher, I. Translating the force-mechano-sensing GPCRs. Am J Physiol Cell Physiol 322, C1047–C1060, doi:10.1152/ajpcell.00465.2021 (2022).

60 Yao, X. et al. Coupling ligand structure to specific conformational switches in the beta2-adrenoceptor. Nat Chem Biol 2, 417–422, doi:10.1038/nchembio801 (2006).

61 Imai, T., Suzuki, M. & Sakano, H. Odorant receptor-derived cAMP signals direct axonal targeting. Science 314, 657–661, doi:10.1126/science.1131794 (2006).

62 Ezan, J. et al. Primary cilium migration depends on G-protein signalling control of subapical cytoskeleton. Nat Cell Biol 15, 1107–1115, doi:10.1038/ncb2819 (2013).

63 Tadenev, A. L. D. et al. GPSM2-GNAI Specifies the Tallest Stereocilia and Defines Hair Bundle Row Identity. Curr Biol 29, 921–934 e924, doi:10.1016/j.cub.2019.01.051 (2019).

64 Kindt, K. S. et al. EMX2-GPR156-Galphai reverses hair cell orientation in mechanosensory epithelia. Nat Commun 12, 2861, doi:10.1038/s41467-021-22997-1 (2021).

65 Reed, R. R. Signaling pathways in odorant detection. Neuron 8, 205–209, doi:10.1016/0896-6273(92)90287-n (1992).

66 Yau, K. W. & Hardie, R. C. Phototransduction motifs and variations. Cell 139, 246–264, doi:10.1016/j.cell.2009.09.029 (2009).

67 Kaupp, U. B. Olfactory signalling in vertebrates and insects: differences and commonalities. Nat Rev Neurosci 11, 188–200, doi:10.1038/nrn2789 (2010).

68 Hoffman, B. U. et al. Focused ultrasound excites action potentials in mammalian peripheral neurons in part through the mechanically gated ion channel PIEZO2. Proc Natl Acad Sci U S A 119, e2115821119, doi:10.1073/pnas.2115821119 (2022).

69 Beaulieu-Laroche, L. et al. TACAN Is an Ion Channel Involved in Sensing Mechanical Pain. Cell 180, 956–967 e917, doi:10.1016/j.cell.2020.01.033 (2020).

70 Pan, B. et al. TMC1 Forms the Pore of Mechanosensory Transduction Channels in Vertebrate Inner Ear Hair Cells. Neuron 99, 736–753 e736, doi:10.1016/j.neuron.2018.07.033 (2018).

71 Burns, J. C., Kelly, M. C., Hoa, M., Morell, R. J. & Kelley, M. W. Single-cell RNA-Seq resolves cellular complexity in sensory organs from the neonatal inner ear. Nat Commun 6, 8557, doi:10.1038/ncomms9557 (2015).

72 Brownell, W. E., Bader, C. R., Bertrand, D. & de Ribaupierre, Y. Evoked mechanical responses of isolated cochlear outer hair cells. Science 227, 194–196, doi:10.1126/science.3966153 (1985).

73 Ashmore, J. Cochlear outer hair cell motility. Physiol Rev 88, 173–210, doi:10.1152/physrev.00044.2006 (2008).

74 Lu, S. et al. Activation pathway of a G protein-coupled receptor uncovers conformational intermediates as targets for allosteric drug design. Nat Commun 12, 4721, doi:10.1038/s41467-021-25020-9 (2021).

75 Liu, Y. et al. Critical role of spectrin in hearing development and deafness. Sci Adv 5, eaav7803, doi:10.1126/sciadv.aav7803 (2019).

76 Nehme, R. et al. Mini-G proteins: Novel tools for studying GPCRs in their active conformation. PLoS One 12, e0175642, doi:10.1371/journal.pone.0175642 (2017).

77 Zhang, Y., Sun, H., Zhang, J., Brasier, A. R. & Zhao, Y. Quantitative Assessment of the Effects of Trypsin Digestion Methods on Affinity Purification-Mass Spectrometry-based Protein-Protein Interaction Analysis. J Proteome Res 16, 3068–3082, doi:10.1021/acs.jproteome.7b00432 (2017).

78 Mastronarde, D. N. Automated electron microscope tomography using robust prediction of specimen movements. J Struct Biol 152, 36–51, doi:10.1016/j.jsb.2005.07.007 (2005).

79 Scheres, S. H. Processing of Structurally Heterogeneous Cryo-EM Data in RELION. Methods Enzymol 579, 125–157, doi:10.1016/bs.mie.2016.04.012 (2016).

80 Zheng, S. Q. et al. MotionCor2: anisotropic correction of beam-induced motion for improved cryo-electron microscopy. Nat Methods 14, 331–332, doi:10.1038/nmeth.4193 (2017).

81 Rohou, A. & Grigorieff, N. CTFFIND4: Fast and accurate defocus estimation from electron micrographs. J Struct Biol 192, 216–221, doi:10.1016/j.jsb.2015.08.008 (2015).

82 Melero, R. et al. Continuous flexibility analysis of SARS-CoV-2 spike prefusion structures. IUCrJ 7, 1059–1069, doi:10.1107/S2052252520012725 (2020).

83 Pettersen, E. F. et al. UCSF Chimera--a visualization system for exploratory research and analysis. J Comput Chem 25, 1605–1612, doi:10.1002/jcc.20084 (2004).

84 Emsley, P. & Cowtan, K. Coot: model-building tools for molecular graphics. Acta Crystallogr D Biol Crystallogr 60, 2126–2132, doi:10.1107/S0907444904019158 (2004).

85 Adams, P. D. et al. PHENIX: a comprehensive Python-based system for macromolecular structure solution. Acta Crystallogr D Biol Crystallogr 66, 213–221, doi:10.1107/S0907444909052925 (2010).

86 Pettersen, E. F. et al. UCSF ChimeraX: Structure visualization for researchers, educators, and developers. Protein Sci 30, 70–82, doi:10.1002/pro.3943 (2021).

